# Aberrant pace of cortical neuron development in brain organoids from patients with 22q11.2 deletion syndrome-associated schizophrenia

**DOI:** 10.1101/2023.10.04.557612

**Authors:** Sneha B. Rao, Zhixiong Sun, Francesco Brundu, Yannan Chen, Yan Sun, Huixiang Zhu, Robert J. Shprintzen, Raju Tomer, Raul Rabadan, Kam W. Leong, Sander Markx, Steven A. Kushner, Bin Xu, Joseph A. Gogos

## Abstract

Adults and children afflicted with the 22q11.2 deletion syndrome (22q11.2DS) exhibit cognitive, social, and emotional impairments, and are at significantly heightened risk for schizophrenia (SCZ). The impact of this deletion on early human brain development, however, has remained unclear. Here we harness organoid models of the developing human cerebral cortex, cultivated from subjects with 22q11.2DS and SCZ, as well as unaffected control samples, to identify cell-type-specific developmental abnormalities arising from this genomic lesion. Leveraging single-cell RNA-sequencing in conjunction with experimental validation, we find that the loss of genes within the 22q11.2 locus leads to a delayed development of cortical neurons. This impaired development was reflected in an increased proportion of actively proliferating neural progenitor cells and a reduced fraction of more mature neurons. Furthermore, we identify perturbed molecular imprints linked to neuronal maturation, observe the presence of sparser neurites, and note a blunted amplitude in glutamate-induced Ca2+ transients. The aberrant transcription program underlying impaired development contains molecular signatures significantly enriched in neuropsychiatric genetic liability. MicroRNA profiling and target gene investigation suggest that microRNA dysregulation due to *DGCR8* deficiency may drive perturbations of genes governing the pace at which maturation unfolds. Using protein-protein interaction network analysis we define complementary effects stemming from other genes residing within the deleted locus. Our study uncovers reproducible neurodevelopmental and molecular alterations resulting from 22q11.2 deletions, with findings that could advance disease modeling and drive the development of therapeutic interventions.

---

Deletions at the chromosome 22q11.2 locus are the most frequent deletion in humans and present a variable clinical phenotype involving multiple systems^1–4^. Adults and children with the 22q11.2 Deletion Syndrome (22q11.2DS) demonstrate cognitive, social and emotional impairments^5,6^. 22q11.2 deletions are also one of the strongest genetic risk factors for schizophrenia (SCZ)^7^. A number of behavioral and neuroanatomical abnormalities in children with 22q11.2DS likely stem from altered early brain development^8–10^. Similarly, for a sizable fraction of patients with SCZ, especially ones with large effect mutations, genetic liability leads to pre-morbid dysfunction before clinical disease onset by disturbing early development^11–14^.

Rare and highly penetrant risk mutations provide an opportunity to leverage experimentally tractable model systems for identifying convergent neural substrates and underlying molecular mechanisms that might serve as entry points to prevent or reverse disease progression. While understanding the full chain of events leading from a risk mutation to the emergence of clinical symptoms necessitates analysis of the intact brain in behaving animals, for mutations associated with neurodevelopmental abnormalities there is a need for such studies to be complemented by human experimental models that mirror prenatal states of the human brain. Forebrain organoid models provide a critical platform to dissect and integrate information, which would be hard to be accessed otherwise in the developing human brain^15–17^.

To this end, here we analyze organoid models of the developing human cerebral cortex derived from hiPSC lines from patients with 22q11.2DS and SCZ as well as unaffected controls matched by sex and age. We emphasize analysis of EN lineage based on plethora of previous studies in animal models indicating that deficits in synaptic functionality of ENs contribute to the cognitive deficits associated with the 22q11.2DS^18–20^. We showed that loss of genes within the 22q11.2 locus leads to neurodevelopmental defects characterized by delayed neuronal maturation trajectories and pervasive changes in gene expression. Transcriptional profiling underscored specific biological pathways underlying the observed abnormal neurodevelopmental trajectory, providing mechanistic insights into the molecular pathogenesis of 22q11.2-associated clinical manifestations and into potential therapeutic interventions. The aberrant transcription program underlying asynchronous development contains molecular signatures significantly enriched in neuropsychiatric genetic liability suggesting that our findings may be generalized to other genetic causes of SCZ, especially ones with large effect mutations. The human forebrain organoid model presented here could serve as a human-specific preclinical model and a resource for further studies of the molecular and cellular pathogenesis of the 22q11.2DS.

## Results

### Forebrain organoids as models of the 22q11.2 deletion

To model and study the consequences of the 22q11.2 deletion in the context of early human neurodevelopment, we derived 3D dorsal forebrain patterned organoids from human induced pluripotent stem cell (hiPSC) lines from patients with the canonical 3-Mb 22q11.2 deletion and SCZ using a well-established protocol^21^ with high reliability of differentiation. For the analysis shown here, we employed iPSC lines from sibling pairs, as well as cases and matched unrelated control pairs from the NIMH Repository and Genomic Resource. Generation of these lines has been described previously^22^ (see also **Methods**). The dorsal forebrain identity and cell type composition of organoids at DIV70 were confirmed through immunostaining using the forebrain marker FOXG1, dorsal forebrain markers PAX6 and TBR1, and layer-specific markers such as PH3 (proliferating NPCs at the SVZ), CTIP2 (lower layers), and SATB2 (upper layers) (**Fig S1a**). Additionally, cell-type-specific markers were used to identify radial glia (Nestin), intermediate progenitor cells (TBR2) (**Fig S1b**), and neurons [somatodendritic marker MAP2 and immature neuron marker doublecortin (DCX)] (**Fig S1c**).

Quantification of the growth rate of patient and control organoids, between DIV14 (Week 2) and DIV140 (Week 10), did not reveal significant genotypic differences in their average area or perimeter [Kolmogorov-Smirnov (KS) test, ns, n=72 control, 76 patient organoids, **Fig 1a,b**]. We also tested whether there are alterations in neuronal activity in patient compared to control organoids using as a proxy system-wide glutamate-induced Ca^2+^ transients. Comparisons of three pairs at DIV 259 identified a significant decrease in Ca^2+^ peak amplitude in case compared to control cells (*P* = 5.78×10^-12^, average number of cells per organoid: n=89 patient, n=124 control, **Fig 1c,d,e**) consistent with previous findings in monolayer cultures of both mouse and patient neurons^22,23,18^. Thus, despite the lack of overt effects in growth, patient organoids display different functional properties, validating their usefulness as a preclinical model for studying the disease mechanisms underlying 22q11.2 deletions.

**Fig 1:**
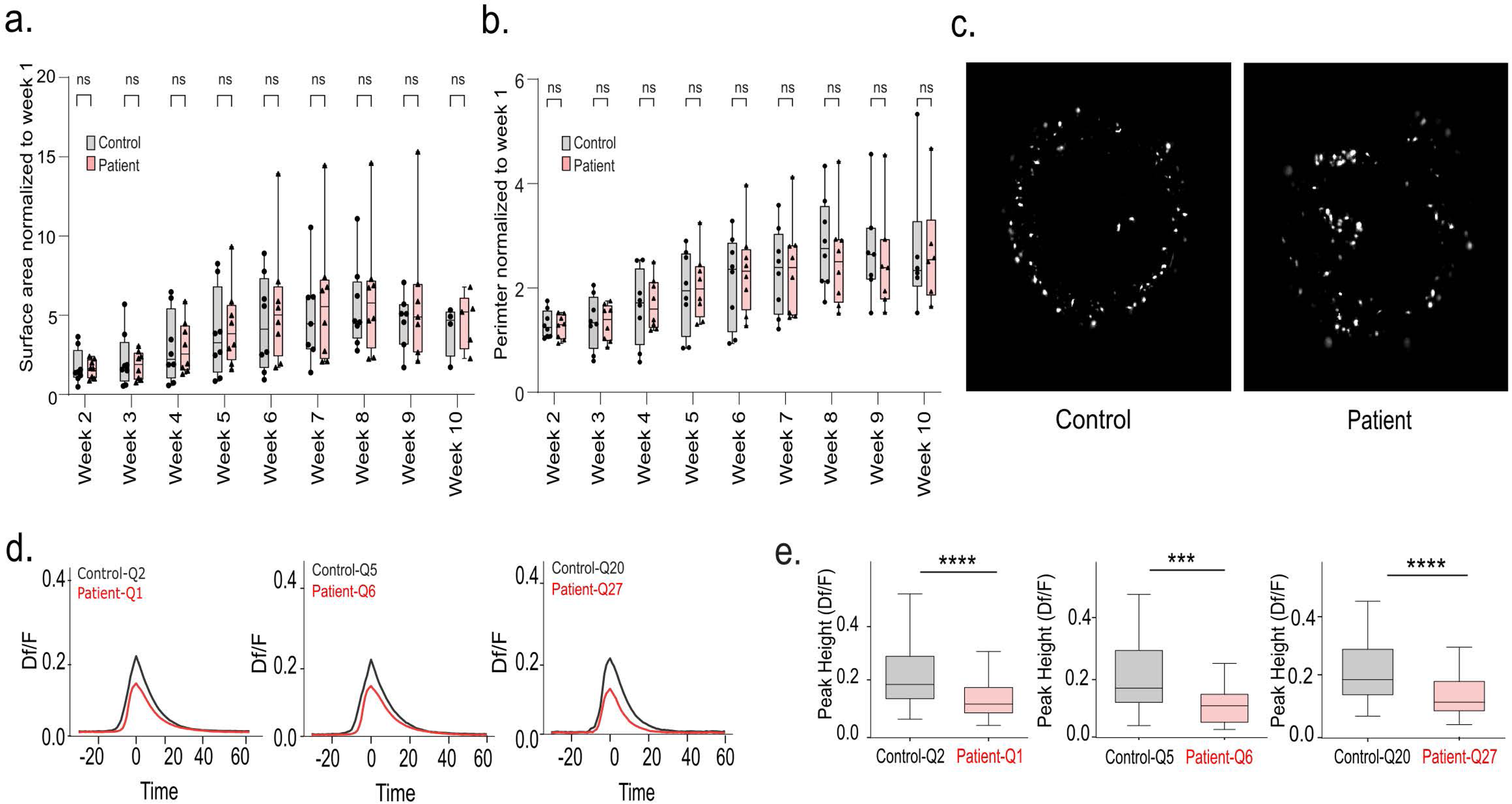
Organoid growth and neuronal activity analysis. **(a)** Box plots depicting increase in organoid surface area in patient and control organoids at different time points normalized to week 1. **(b)** Box plots depicting increase in organoid perimeter in patient and control organoids at different time points normalized to week 1. **(c)** Representative denoised and segmented epifluorescence microscope images (4x) of Ca^2+^ unit signals detected in organoids *in situ* **(d)** Average peak-aligned traces (DF/F vs time, zero corresponding to peak time) of Ca^2+^ transients detected in different units (cells) in patient (red) and control (black) organoids. Three patient and three control organoids from two independent batches analyzed at this timepoint (average number of cells per organoid: n for patient = 89, n for control = 124). *Left:* Combined aligned traces from batches 1 and 2, *Middle:* Batch 1, *Right:* Batch 2. **(e)** Quantification of Ca^2+^ peak amplitude in patient (red) and control (black) organoids. Patient organoids show significantly lower peak amplitude. *Left:* Combined batches 1 and 2 (KS test, *P* = 5.78×10^-12^), *middle:* Batch 1 (KS test, *P* = 3.90×10^-3)^, *right:* Batch 2 (KS test, *P* = 3.77×10^-10^).

To identify early neurodevelopmental disease pathways leading to disease phenotypes, dissect in more detail the various cell population alterations and directly examine genotypic differences in transcriptomic and regulatory profiles during neurogenesis and neuronal differentiation, we used single cell RNA sequencing (scRNA-seq). We analyzed the cell type composition and molecular signatures of patient and control forebrain organoids at two key stages: DIV70, characterized by abundant progenitors and the onset of robust neurogenesis, and DIV150, marked by the emergence of diverse cortical excitatory neuron (EN) subtypes, interneurons, and astroglia. We generated a high viability (>90%) single cell suspension for scRNA-seq from each of 3 pairs of DIV70 and DIV150 organoids, a total of 6 lines from 3 individual patients and 3 control with 3 organoids per line and developmental time points. To ensure consistency, data from all samples and both time points were first aggregated using CellRanger and processed through a uniform and harmonized analysis pipeline. Quality control metrics were consistent across organoids from all iPSC lines at both time points, showing low detection of mitochondrial and ribosomal genes, with comparable count metrics across all samples (**Fig S2a**). We observed negligible differences in the expression of hypoxia and apoptosis markers between patient and control organoids and between time points, indicating healthy organoids across time points and genotypes (**Fig S2b, c**). Following quality control and filtering, a total of 55,762 single cells were analyzed. We identified 8 distinct cell clusters using the Leiden algorithm, with a uniform distribution across all samples, indicating consistent cell composition among the samples^24,25^ (**Fig S2d**). Assessment of 22q11.2 locus genes expressed in forebrain organoids confirmed the expected ∼50% reduction in all patients compared to control organoids (**Fig S2e**), indicating high consistency of the transcriptomes (**Fig S2f**).

### Altered cell type proportions in patient organoids

To more precisely annotate cell types observed in the scRNA-seq datasets, we used as a reference a published dataset that classifies the signature transcriptomic profiles of cell types in the developing human telencephalon at various stages of development^26^. Cell type mapping was conducted using the singleR tool^27^. By using a custom-built stratified subsampling approach (see Methods) we confirmed that 98% of cell annotations passed the threshold with high confidence. We found that the majority of cell types found in the developing human brain are represented in our DIV70 and DIV150 datasets (**Fig 2a, Fig S3a and b**). We further confirmed the specificity of cell type annotation using conventional cell type specific markers **(Fig S3 c and d**). At DIV70 our annotation strategy identified early radial glia (RG-early) and ENs as the predominant cell types. Intermediate progenitors differentiating into newborn ENs (IPC-nEN) represent the next most abundant group. By DIV150 organoids display a diversity of cell types including EN, late radial glia (RG-late), cortical as well as striatal interneurons (IN) and astrocytes. Overall, as development progresses, the fraction of EN lineage cells (defined by RG-early, RG-div, IPC-div, IPC, nEN and EN) is altered (**Fig S3e**). Importantly, while the relative proportion of EN lineage progenitors decreases from DIV70 to 150, the proportion of newborn and maturing EN remains fairly constant. Conversely, there is an expansion of IN lineage as well as astrocytes at DIV150 (**Fig 2a, b**).

**Fig 2:**
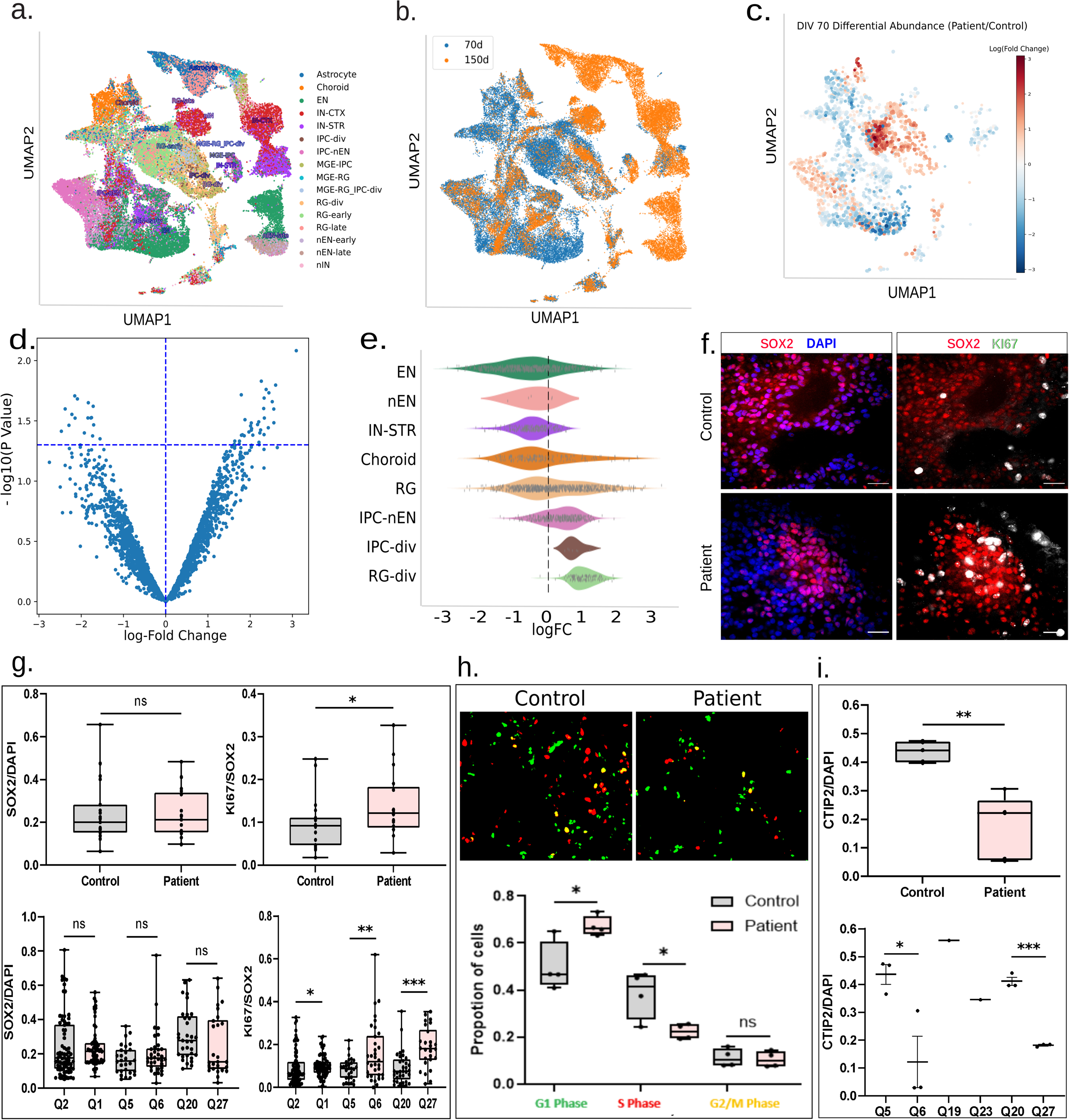
Cell type composition and proliferation analysis of organoids. **(a)** Annotation of cell types in the dataset with reference to human fetal brain^26^. Most of cell types in the developing human brain are successfully recapitulated. (b) UMAP color-coded to highlight distribution of annotated cell types in DIV70 (blue) and DIV150 (orange). (c) Density maps depicting the differential abundance (log fold changes) in all cell types between patient and control organoids at DIV70. There is a higher cell density in progenitor regions (RG-div, IPC-div, IPC-nEN) and lower cell density in differentiated neuron regions (EN) in patient organoids compared to control. **(d)** Volcano plot illustrating differentially abundant neighborhoods between patient and control groups using Milo. The y-axis represents – log_₁₀_(spatial FDR), and the x-axis shows the log_₂_fold change (patient/control). The horizontal blue dotted line indicates the FDR = 0.05 threshold **(e)** Beeswarm plot visualizing the distribution of log fold changes across neighborhood annotations for each cell type in DIV70 organoids between patient and control groups, analyzed using Milo software. The dashed line represents no change. **(f)** Representative confocal microscope images (40x) of a single 2D Z-stack of control (*top*) and patient (*bottom*) organoid whole-mount preparations stained for cell cycle marker, Ki67 (white). Depicted are DIV70 organoids double labelled for NPC marker SOX2 (red) and nuclear marker DAPI (blue) (*left*), NPC marker SOX2 and active cell cycle marker Ki67 (*right*). Scale bar = 20µm. **(g)** Quantification of the fraction of cells labelled with the aforementioned markers reveals that the fraction of active cycling progenitors is significantly higher in patient organoids (n for patient = 17, n for control = 19, from 3 independent batches). Total number of SOX2 positive NPCs as percentage of DAPI-labeled cells (upper left panel); Fraction of active cycling (Ki67-positive) NPCs (SOX2-positive) (KS test, *P* = 0.04) (upper right panel). Number of SOX2-positive NPCs represented as percentage of DAPI-labelled nuclei in individual patient-control pairs (KS test, *P* = 0.01) (lower left panel); Fraction of active cycling (Ki67-positive) NPCs (SOX2-positive) (KS test, *Ppair1* = 0.25, *Ppair2* = 0.01, *Ppair3* = 5.78×10^-5^) (lower right panel). **(h)**. Representative images of PIP-FUCCI-labeled NPCs in controls and patients (upper panel). Quantification of the fraction of NPCs in G1, S, and G2/M phases reveals a significant accumulation of G1 phase NPCs in patient samples (two-sided t-test *P* = 0.045, fold change = 1.37) and lower number of NPCs in the S phase (two-sided t-test *P* = 0.035, fold change = 0.58) (lower panel). **(i)**. Quantification indicates a significantly lower fraction of NPC-derived neurons in patients compared to controls (KS test, *P* = 0.01). The fraction of neurons (CTIP2-positive) is shown as a percentage of DAPI-labeled cells (upper panel). The number of CTIP2-positive neurons, expressed as a percentage of DAPI-labeled nuclei for individual patient-control pairs, is presented (lower panel). Control samples include Q2, Q5, QR19, and QR20; patient samples include Q1, Q6, QR23, and QR27.

We used Milo^28^, a scalable statistical framework for differential abundance testing, to analyze cell type distributions in patient organoids. Quantification of various cell types revealed a nominally significant higher abundance of cycling radial glial (RG-div) and cycling intermediate progenitor cells (IPC-div) alongside a lower abundance of differentiated ENs in patient organoids compared to control organoids (**Fig 2c, d, e**). To further validate changes in cycling NPCs, we performed 3D immunostaining of DIV70 organoids. We used Ki67, a proliferation marker, to label actively cycling cells, and SOX2, to identify NPCs (**Fig 2f**). Consistent with the scRNA-seq results, our analysis revealed a significant increase in the proportion of Ki67-positive/SOX2-positive cells (cycling NPCs) in patient organoids compared to controls (**Fig 2g, upper panel right**). There was a consistency across each line of the control and patient organoids, demonstrating reasonable reproducibility among organoids within each group (**Fig 2g, lower panel**). However, the total number of NPCs (SOX2-positive) remained unchanged (**Fig 2g, upper panel left**).

To further investigate the proliferation and neurogenesis characteristics of cycling NPCs, we performed two additional analyses. First, purified NPCs (SOX2+) were isolated from DIV20 organoids and cultured in 2D. These cells were transduced with the PIP-FUCCI biosensor (**Fig. 2h, upper panel**), and the proportion of cells in each cell cycle phase (G1, S, G2/M) was quantified. The results showed a significantly higher percentage of cells (t-test *P* = 0.037, fold change = 1.35) in the G1 phase and a significantly lower percentage (t-test *P* = 0.045, fold change = 0.58) in the S phase in patient NPCs compared to controls (**Fig. 2h, lower panel**). Second, we used fluorescence-activated cell sorting (FACS) to experimentally validate the reduction in neuron numbers suggested by the scRNA-seq analysis. Purified NPCs (SOX2+) were isolated from all DIV20 organoids to form 3D neurospheres, which were then cultured in neuronal differentiation medium for 7 days before being dissociated into single cells. The cells were fixed and immunostained with SOX2 and CTIP2, and the proportions of each cell type in the neurospheres were analyzed using FACS. Initially, SOX2+ cells constituted the majority (∼90%) of the total population in neurospheres. After 7 days of differentiation, CTIP2+ neurons increased to ∼40% of the total population in control-derived neurospheres. However, patient-derived neurospheres showed a marked reduction in CTIP2+ neurons (∼20%) (**Fig. 2i**). These differences were statistically significant (t-test *P* < 0.01) and consistent across patient-control pairs (**Fig. 2i**). A parallel assay assessing layer thickness in DIV45 organoids, where rosette structures are prominent, yielded consistent results. The PAX6-positive NPC layer was defined as the subventricular zone (SVZ), while the CTIP2-positive neuron layer represented the cortical plate (CP). Although no significant change was observed in the relative thickness of the SVZ, consistent with a total NPC count in patient organoids comparable to controls, a significant decrease in CP thickness was detected in patient organoids (**Fig. S4a-c**). Collectively, these findings suggest that the 22q11.2 deletion delays NPC differentiation during early neurodevelopment.

### 22q11.2 deletion leads to asynchronous developmental trajectories in forebrain organoids

To determine whether differences in cell population frequencies arise from aberrant developmental trajectories during early neuronal development, we used Monocle3^29^ to examine cell distribution across pseudotime in a cell type-specific manner (**Fig S5**). This analysis focused on the developmental dynamics of the affected cell types within the EN lineage in our scRNA-seq datasets (**Fig S5a,b**). Plotting the cell distribution of DIV70 EN lineage along the pseudotime trajectory revealed an increased distribution of patient cells in the early phase of the trajectory (progenitor states) and a decreased distribution of patient cells towards the terminal phase of the trajectory (mature neuron states) (*P* = 1.347×10^-11^, two-sided KS test; **Fig 3a, b**). These findings indicate delayed differentiation of progenitors within the EN lineage in patient organoids.

**Fig 3:**
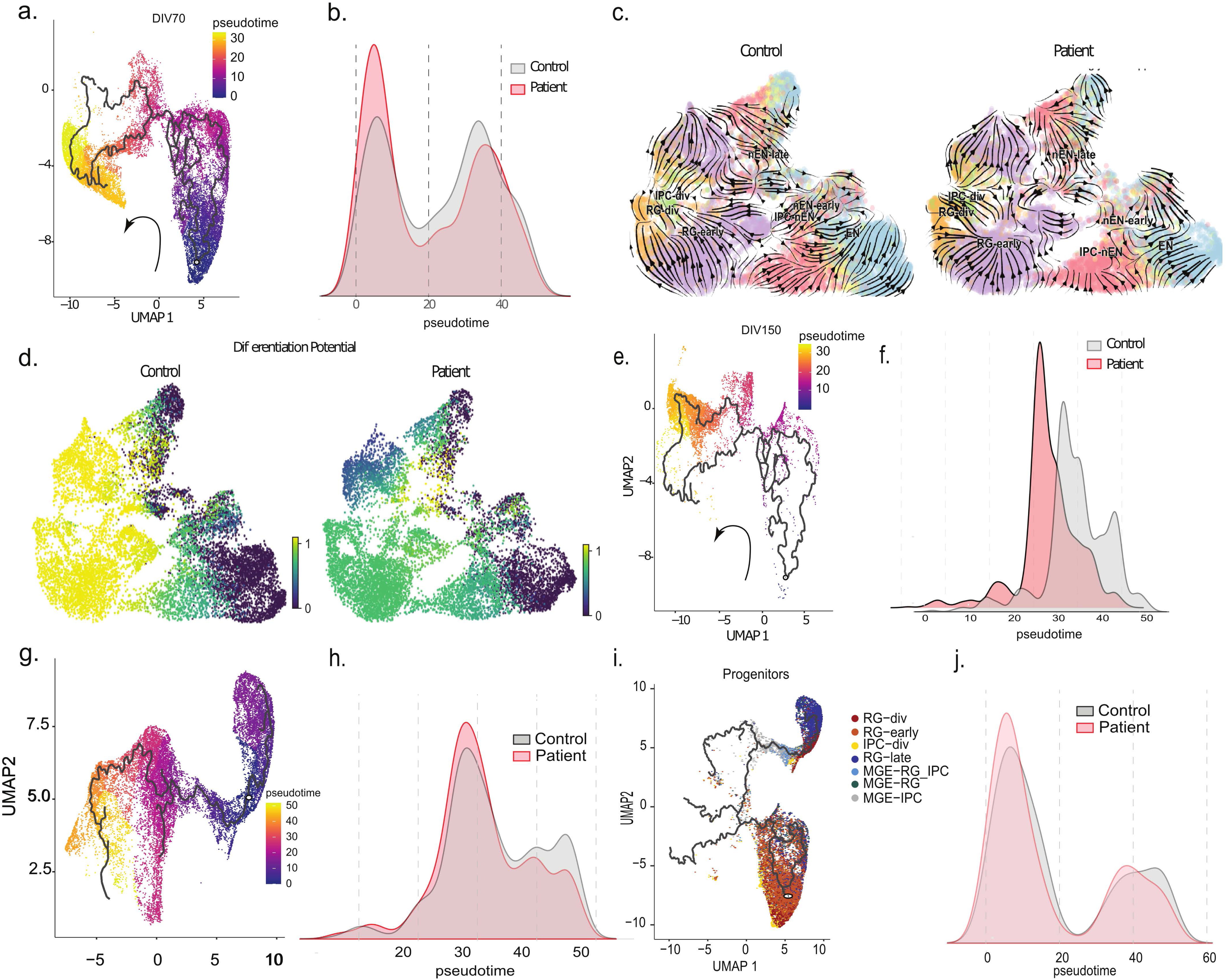
Pseudotime analyses of the pace of neuronal development. **(a)** Pseudotime trajectory of the EN lineage at DIV70 colored by pseudotime. **(b)** Ridge plot depicting distribution of cells in patient and control EN lineage at DIV70. Patient cell distribution is shifted towards early points of the trajectory represented by progenitors while control cell distribution is more dispersed. There is a significant difference between the two distributions (KS test, *P* = 1.33×10^-11^). **(c)** RNA velocity embedding stream at DIV70 depicting the EN lineage trajectory in control (*left*) and patient (*right*) organoids. **(d)** UMAP plots colored by differentiation potential in control (*left*) and patient (*right*) DIV70 EN lineage. Progenitors in control organoids show high differentiation potential while progenitors in patient organoids show low differentiation potential. **(e)** Pseudotime trajectory at DIV150 colored by pseudotime. **(f)** Ridge plot showing distribution of cells in patient and control DIV150 EN lineage. Patient cell distribution is largely shifted to early points of the trajectory represented by more immature stages of the lineage while control cell distribution is shifted towards later points along the pseudotime. There is a significant difference between the two distribution (KS test, *P* = 2.13×10^-4^). **(g)** Pseudotime trajectory of cells in patient and control IN lineage colored by pseudotime. **(h)** Ridge plot depicting distribution of cells in patient and control IN lineage along pseudotime. Patient cell distribution is shifted towards early points along pseudotime while control cell distribution is more dispersed. There is a significant difference between the two distributions (KS test, *P* = 1.33×10^-2^). **(i)** Pseudotime trajectory of progenitor populations generated with Monocle3. Trajectory starts at RG-div and cyc-IPC and progresses to RG-early. Later pseudotimes are represented by RG-late and IN progenitors such as MGE-RG, MGE-IPC and MGE-RG-IPC-DIV. **(j)** Ridge plot depicting distribution of progenitors in patient and control lineage Patient cell distribution is shifted largely to early points of the trajectory represented by earlier progenitors while control cell distribution is shifted more towards later time points. There is a significant difference between the two distribution (KS test, *P* = 4.53×10^-6^), indicating a developmental delay at the level of the progenitors.

To validate this observation using a complementary approach, we applied RNA velocity analysis with scVelo to infer directed developmental dynamics from our single-cell transcriptomics dataset. RNA velocity analysis uses gene splicing dynamics to quantitatively measure transcriptional dynamics of each gene. Gene velocity information was used by CellRank^30^ to define cell transition states and infer the differentiation trajectory and fate potential of individual cells. We used the CellRank pipeline to order cells based on latent time as determined by the transition of RNA velocity between cell states and computed cell-to-cell transition probabilities using a Markov state model of RNA velocity and transcriptomic similarity. We defined EN as the terminal state and computed absorption probability (a measure of the probability of each cell reaching the endpoint of the lineage) and differentiation potential (a measure of the entropy around the absorption probabilities for each cell and its potential to differentiate towards the terminal state). UMAP plot of the velocity stream, generated by combining the Cellrank probabilities and latent time from scVelo^31^, revealed a difference in the velocity, especially in the IPC-nEN cluster (**Fig 3c**). Computation of Differentiation Potential in DIV70 batch-corrected datasets indicated that cells in the EN lineage tend to stay in the progenitor status in patient compared to control organoids (**Fig 3d**). Taken together, these results suggest that the probability of RG or IPC cells reaching the terminal EN state in patient organoids is lower than in control.

A significant lag in the patient EN development with fewer patient neurons towards the terminal portion of the pseudotime trajectory was still detectable at DIV150 (*P* = 2.13×10^-4^, two-sided KS test) indicating that delayed maturation of EN persists at later developmental stages (**Fig 3e,f**). Pseudotime analysis in the IN lineage at DIV 150 similarly showed a reduced distribution of IN towards the endpoint of the developmental trajectory (*P* = 0.013, two-sided KS test) indicating a significant delay in the differentiation of IN as well (**Fig 3g,h and Fig S5c**). Convergent effects on two neuronal lineages suggest that the 22q11.2 deletion may affect developmental timelines by acting at an earlier common progenitor stage. A delay in the differentiation of the RG-early for example, could in effect delay EN specification, but also the specification of RG-late and MGE progenitors. To test this hypothesis, we sub-grouped progenitors and examined their distribution along pseudotime in the combined DIV70 and DIV150 datasets. This analysis showed a significant shift in the distribution of cells towards earlier progenitor states along a trajectory progressing from early (RG-early) and cycling RG (RG-div) as well as dividing intermediate progenitor cells to late RG (RG-late), MGE-RG and MGE-IPC in patient organoids (*P* = 4.53×10^-6^, two-sided KS test) (**Fig 3i,j**). Thus, a defect traced back to early progenitors may impact the development trajectory of both EN and IN lineage progenitors.

### Patient EN display impaired maturation signatures

The finding that NPCs are more likely to remain in a cycling state and fail to differentiate efficiently into neurons in patient organoids prompted us to determine whether those patient neurons that do differentiate show further impairments in their growth, by looking for impaired maturation signatures in morphological assays as well as in bulk RNA-seq. We transduced patient and control organoids with an AAV-hSYN1-eGFP virus at 50-55 days and 2 weeks later we imaged them as a whole mount for differences in neurite outgrowth. This strategy facilitated imaging of a large area of the organoid without sectioning, thus preserving 3D cell morphology (**Fig 4a**). Immunohistochemical analysis on these neurons showed that they express the somatodendritic marker MAP2 and the immature neuron marker DCX (**Fig S1c**). We quantified the number of branchpoints, number of primary neurites and total neurite length for each traced cell. We analyzed a total of 8-11 organoids per line over 3-4 separate batches generated on different dates. A total of 66-166 cells per iPSC line were analyzed. We observed a marked reduction in the neurite complexity of the patient organoid neurons compared to control, with a significant reduction in all neurite complexity indices (**Fig 4b**). Importantly, these differences were reliably observed in each of the individual patient/control pairs (all three pairs analyzed above and one additional case and matched unrelated control pair, **Fig 4b bottom panel**). A combined analysis revealed that the reduction in neurite complexity in patient organoids was highly significant with respect to number of branchpoints (t-test, *P* = 0.013, **Fig 4b**, upper left), neurite length (t-test, *P* = 0.0209, **Fig 4b**, upper middle) and number of primary neurites (t-test, *P* = 0.0038, **Fig 4b**, upper right). In sum, patient neurons during early forebrain development present stunted neurite growth indicative of delayed maturation.

**Fig 4:**
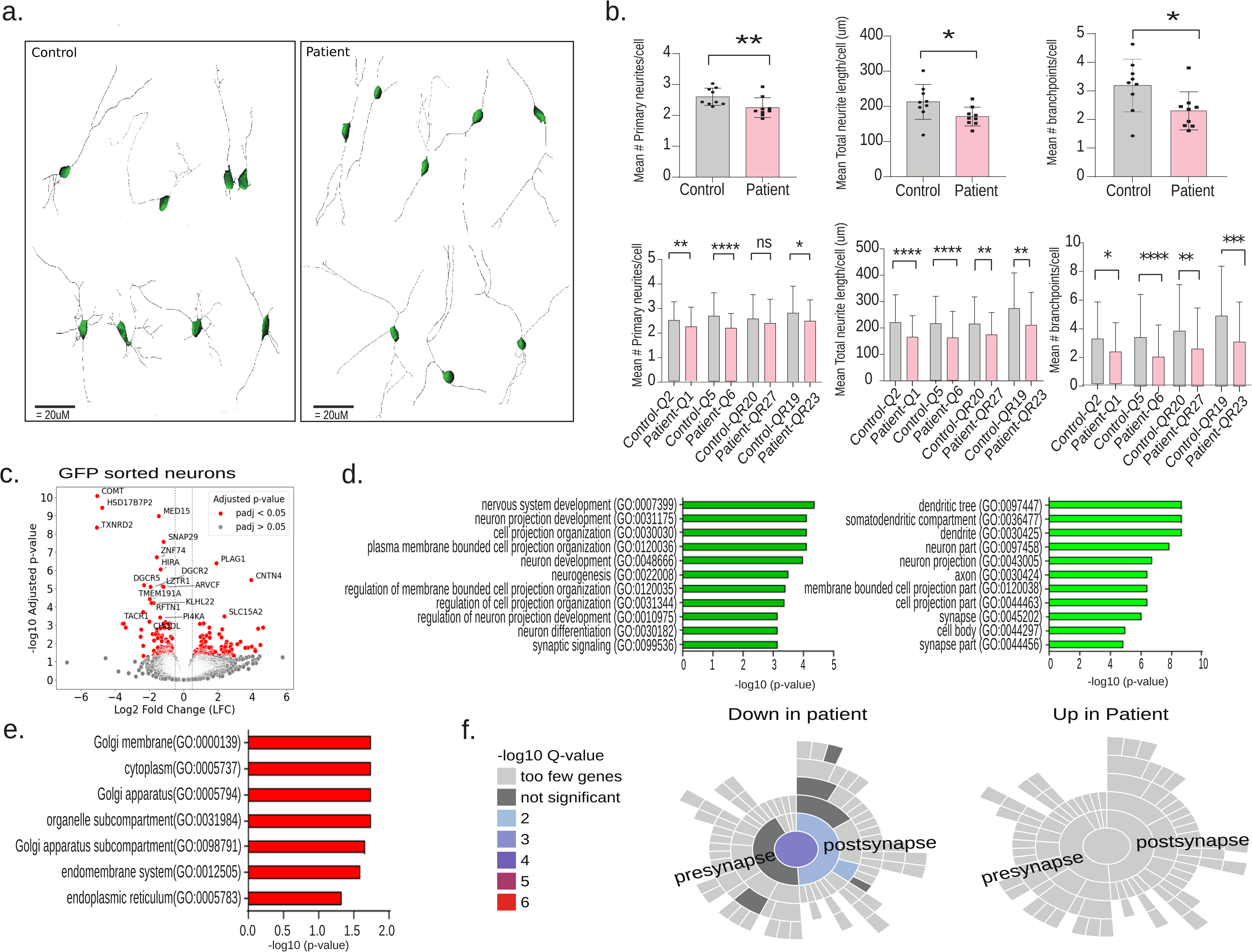
Morphological and transcriptional signatures of impaired EN maturation. **(a)** Representative 3D traces of neurons generated from confocal images of DIV70 cultures of control (*left*) and patient (*right*) organoids. **(b)** Quantification of combined patient and control number of primary neurites per cell (*upper left*), total neurite length (*upper middle*) and number of branchpoints per cell (*upper right*) across multiple experiments (unpaired two-sided t-test, *P* < 0.05). 8-11 organoids were analyzed per line over 3-4 batches for a total of 66-166 cells for each line. Data are presented as mean ± SD, each point represents averaged data from a unique line and batch. Quantification of number of primary neurites per cell (*bottom left*), total neurite length (*bottom middle*) and number of branchpoints per cell (*bottom right*) for different patient and control paired organoids. Data are presented as mean ± SD. Across all pairs, patient neurons have a significantly simpler neurite architecture (two-sided Mann Whitney Test, *P* < 0.05). Neurons analyzed with immunostaining were positive for MAP2 and immature neuron marker DCX (see also **Fig S1b**, *bottom*). **(c)** Volcano plot showing differentially expressed genes in sorted neurons from DIV70 patient organoids compared to control. **(d,e)** GO term enrichment analysis of top downregulated **(d)** and upregulated **(g)** DEGs in patient neurons. **(f)** SynGO gene enrichment analysis of genes downregulated (*left*) and upregulated (*right*) in patient neurons. There is a significant enrichment of synaptic genes among the downregulated genes (*P* = 0.00005), with a particular enrichment for transcripts with presynaptic/synaptic vesicle functions (GO:0045202/GO:0098793, *P* = 0.0331), but no significant enrichment of synaptic genes among upregulated genes.

In parallel assays, we used the same set of patient and control organoids (8-10 organoids per line, 4 patient and 4 control lines) transduced with the AAV-hSYN1-GFP virus to isolate GFP-positive neurons using FACS and conduct bulk RNA sequencing on sorted neurons for molecular signatures consistent with delayed maturation. Principal component analysis (PCA) revealed that genetic ancestry (PC1, 35.7%) and genotype (PC2, 23.7%) were the primary sources of transcriptional variance among cortical neurons (**Fig S6**). Differential expression analysis identified 241 genes (143 downregulated and 98 genes upregulated) that were differentially expressed between patient and control neurons (*Padj* < 0.05, |log2(fold change)| > 0.5, **Fig 4c**).

We proceeded to annotate the function of the differentially expressed genes (DEGs) performing Gene Ontology (GO) enrichment analysis. Of the downregulated genes identified in patient neurons, the GO terms were predominantly related to neuron projection development and differentiation as well as to protein localization in dendrites, axons and synapses (**Fig 4d**) (**Supplemental Table 1**). Among the upregulated genes identified in patient neurons, the GO terms enriched were related to the secretory pathway, with protein localization in endoplasmic reticulum and Golgi apparatus (**Fig 4e**) (**Supplemental Table 1**). We also employed synaptic gene ontologies to search for signatures of impaired maturation of synaptic functional pathways among DEGs using the SynGO tool^32^. We found significant enrichment of synaptic genes among the downregulated genes [22 of the 143 transcripts with decreased abundance in patient neurons possessed a synaptic process annotation in SynGO (*P* = 5×10^-5^), with a particular enrichment for transcripts with presynaptic/synaptic vesicle functions (GO:0045202/GO:0098793, *P* = 0.0331), but no significant enrichment of synaptic genes among upregulated genes (**Fig 4f**). The downregulation of synaptic genes may indicate a delay in synapse maturation in patient neurons. Although the challenge of identifying clear synaptic structures in brain organoids limits our ability to directly observe and confirm these abnormalities at the morphological level, the identified molecular signatures support our observation of curtailed development in patient ENs and reinforce the notion that progenitors in the developing patient forebrain fail to differentiate into neurons in a timely manner

### Aberrant gene expression underlying altered EN developmental trajectories

We used the MAST differential expression analysis^33^ to identify DEGs across the EN lineage that may contribute to the aberrant unfolding of neuronal development. Gene expression profiling and functional enrichment analysis identified several genes and genetic pathways important for brain development and function that are altered in patient compared to controls, across various EN developmental stages, clearly indicating that the pace of early embryonic neuronal development of patient organoids is not properly orchestrated at the gene expression level (**Fig 5 and Supplemental Tables 2, 3**). For example, genes upregulated in patient DIV70 RG-div (such as *CCNF, KIF20A, KIFC1, TUBA1C, KIF18B, TOP2A, CKAP2, CDCA2, CCND2, CDC20*), are primarily involved in the regulation of the cell cycle and promotion of cell division whereas downregulated genes (such as *PTN, CDC42, STMN4, EFNB1, EMX2, VEGFA, MAP4, SOX6, POU3F2, PAK3*) are involved in neurogenesis and neuronal differentiation, underscoring how, already at early stages of development, 22q11.2 locus haploinsufficiency alters the transcriptional regulation of progenitor proliferation and differentiation (**Fig 5a,b** and **Supplemental Tables 2, 3**). By contrast, genes upregulated in patient DIV70 EN are significantly enriched for functions related to negative regulation of neurogenesis and neuron growth and guidance pathways (such as *EPHA4, MAP4K4, HES4, LHX2, NTRK3, SEMA6A*) while downregulated genes are primarily involved in neuronal differentiation and maturation pathways (such as *PAK3, RBFOX2, NRP2, FZD3, ANK3, CALR, RTN4*) (**Fig 5c,d** and **Supplemental Tables 2, 3**). Furthermore, as differentiation unfolds in DIV150, patient EN display consistent signatures of impaired maturation as reflected by a lower abundance of transcripts related to neuron migration, maturation and synaptic function (such as *SEMA6A, DAB1, COUPTF-1, SYN10, SYNPR*) and higher abundance of transcripts that are typically produced during periods of active neuronal growth (such as *STMN2, CNTN1, MAPT*) (**Supplemental Tables 4, 5**).

**Fig 5:**
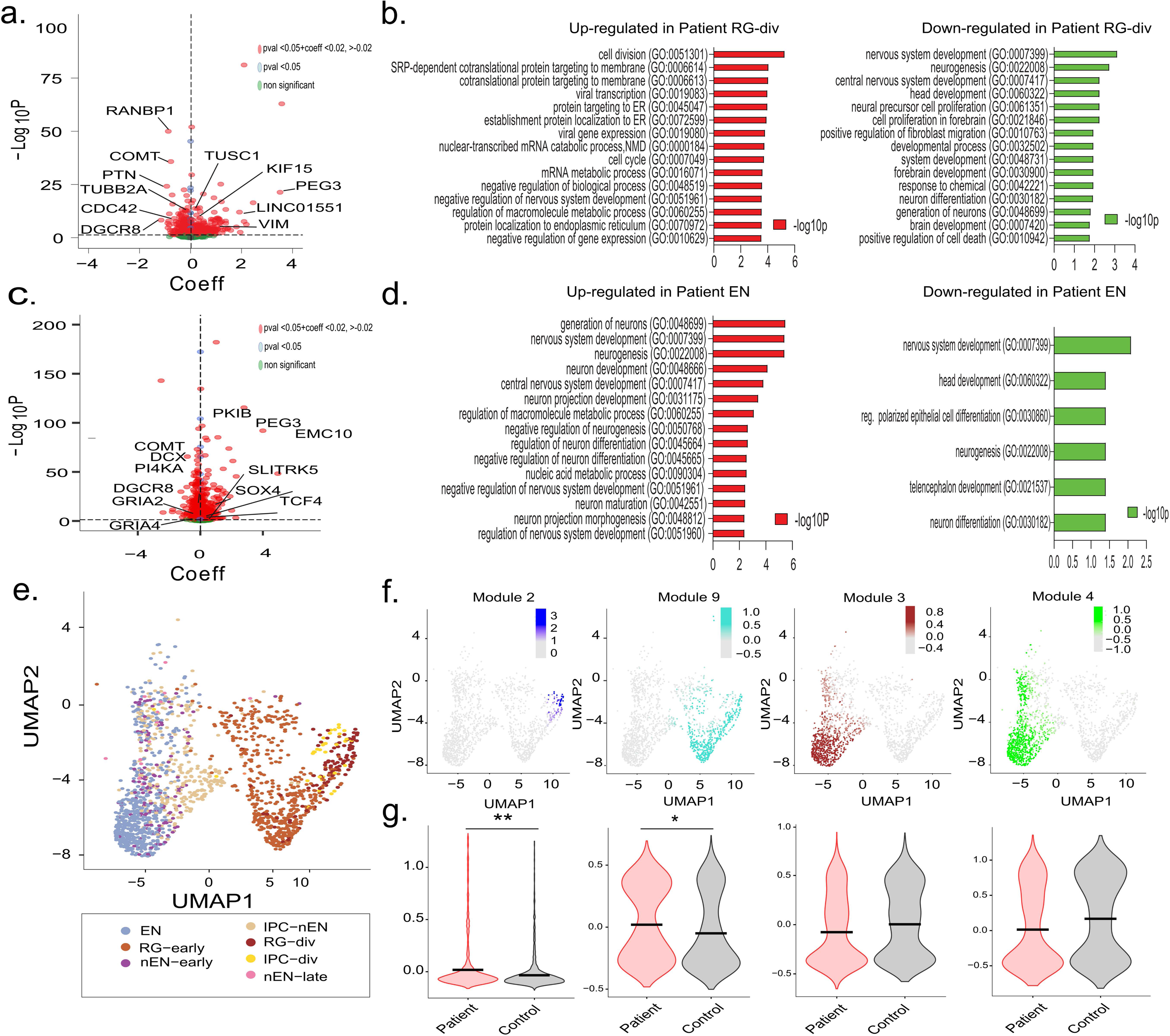
Gene modules and functions dysregulated in the EN lineage of patient organoids. **(a)** A volcano plot displaying DEGs identified in patient-derived RG-div compared to control is shown, with the x-axis representing the log odds of expression between genotypes (Coeff). Dotted lines indicate the thresholds for Coeff > 0.02 or < -0.02. The gene symbols for the top 15 DEGs are labeled. **(b)** GO term enrichment analysis of DEGs in patient RG-div. Cell cycle and proliferation genes are upregulated while neurogenic genes are downregulated. **(c)** A volcano plot displaying DEGs identified in patient-derived ENs compared to control is shown, with the x-axis representing the log odds of expression between genotypes (Coefficient, Coeff). Dotted lines indicate the thresholds for Coeff > 0.02 or < -0.02. The gene symbols for the top 15 DEGs are labeled. **(d)** GO term enrichment analysis of DEGs in patient ENs. Genes related to negative regulation of neurogenesis are upregulated while genes related to promotion of neurogenesis are downregulated. **(e)** UMAP of cell clusters in EN lineage cells in control organoids. **(f)** Identification of clusters of co-expressed genes using scWGCNA in control EN lineage cells. From *left* to *right*: Modules 2 and 9 are expressed in cycling progenitors and progenitor cells respectively and modules 3 and 4 are expressed in EN clusters. **(g)** Scanpy scoring of module genes identified in **(e)** and **(f)** in control and patient EN lineage cells. Expression of modules 2 and 9 are significantly lower (module 2, *P* = 0.002591; module 9, *P* = 0.03008) in control compared to patient cells.

Similar results were obtained by analysis of transcriptional regulatory networks using the pySCENIC pipeline^34^. Transcription factor (TF) regulons with lower activity in patient RG-div are related to nervous system development and neurogenesis. In the EN cluster, TF regulons with lower activity in patient organoids are related to more mature stages of neuron development, neurite growth and synaptic function (such as *SNAP25, STMN2, NRXN1, MAP1B, ANK3, MAPT, CRMP1*), while regulons with higher activity include TFs that are typically abundant and active during early neuronal development stages (such as *LHX2, NEUROD2, SOX4, SOX11, TCF4, FOXG1, POU3F3*) (**Supplemental Table 6**).

To obtain a more unbiased evaluation of the genetic ensembles underlying the observed digression from proper developmental trajectories we performed co-expression analysis on EN lineage cells from DIV70 control organoids using a scWGCNA pipeline^35^. We identified 10 distinct WGCNA modules. Based on the number of cells expressing the modules and module size, we narrowed down our analyses to 4 modules (**Fig 5e, f**) (**Supplemental Table 7**, modules 2, 3, 4 and 9). The largest modules (modules 2 and 9) contain multiple genes that are highly expressed in NPCs and are associated with cell cycle regulation. Genes in modules 3 and 4 are highly expressed in differentiated EN and are related to neuron development, neurite outgrowth and cytoskeleton dynamics (**Fig 5f** and **Supplemental Table 7**). Comparative scWGNCA showed that these modules are preserved in the patient EN lineage but may have differences in expression, density and connectivity. We computed a module score for these gene sets based on their expression level within the EN lineage cells in case and control to indicate if the module is up or downregulated. We further fitted the scores to a linear model with genotype as the predictor, each individual line and batch as the random variables. We found that both progenitor modules (2 and 9) were significantly upregulated (*P* = 0.002 for module 2 and *P* = 0.03 for module 9) in patient compared to control organoids (**Fig 5g**), further supporting our findings that progenitors in patient organoids are more likely to remain in a cycling state. Consistent with pseudotime progression, neuronal modules 3 and 4 were downregulated in patient organoids compared to control, although the difference in the module score was not significant. Interestingly ∼21% and 11% of genes in modules 3 and 4 respectively overlap with the DEGs detected in EN. Similarly, ∼22% and ∼34% of genes in modules 2 and 9 overlap with the DEGs detected in RG-div suggesting that a substantial portion of DEGs contribute to the key co-expression modules in these cell types. In sum, a sizable portion of the genes in each of these control WGCNA modules are dysregulated in patient organoids, indicating a marked dysregulation in the expression of crucial gene expression networks involved in neuronal maturation due to the 22q11.2 deletion. This data provides further evidence that ensembles of genes important for neuron development are dysregulated in progenitor cells of patient organoids, in a pattern consistent with our observation that 22q11.2 deletions impede the pace of early embryonic neuronal development.

### 22q11.2-associated genes and genetic pathways underlying altered EN development

We further evaluated the genes and genetic pathways within the 22q11.2 locus that contribute to the altered early neurodevelopmental trajectories in patient organoids. We first examined the dysregulation of microRNAs (miRNAs) as a potential mechanism for the asynchronous expression of maturation-associated genes. 22q11.2 deletion results in brain-enriched miRNA downregulation and an increase in target gene expression due to (i) hemizygosity of *DGCR8*^36^, a component of the “microprocessor” complex that is essential for miRNA production and (ii) hemizygosity of miRNA genes residing within the deletion^22,20^. To assess the contribution of miRNA dysregulation to our observed cellular and RNA-seq phenotypes we performed bulk miRNA sequencing on a pool of 8-10 DIV70 organoids from 3 patients and 3 controls. We identified 19 miRNAs that were significantly differentially expressed in patient vs control organoids (**Fig 6a**) (**Supplemental Table 8**). Of these, 15 miRNAs were significantly downregulated in patients (*Padj* < 0.05) (**Fig 6a**) (**Supplemental Table 8**), including *hsmir-185* that resides in the 22q11.2 locus. We used the mirNet 2.0 tool^37^ to perform target enrichment and network analysis for these downregulated miRNAs and found 10 miRNA target clusters (**Fig 6b**). GO term enrichment analysis on this target interaction network identified categories encompassing cell cycle-related functions (**Fig 6c**), suggesting a contribution of altered miRNA signaling to the aberrant cell cycle programming in patients.

**Fig 6:**
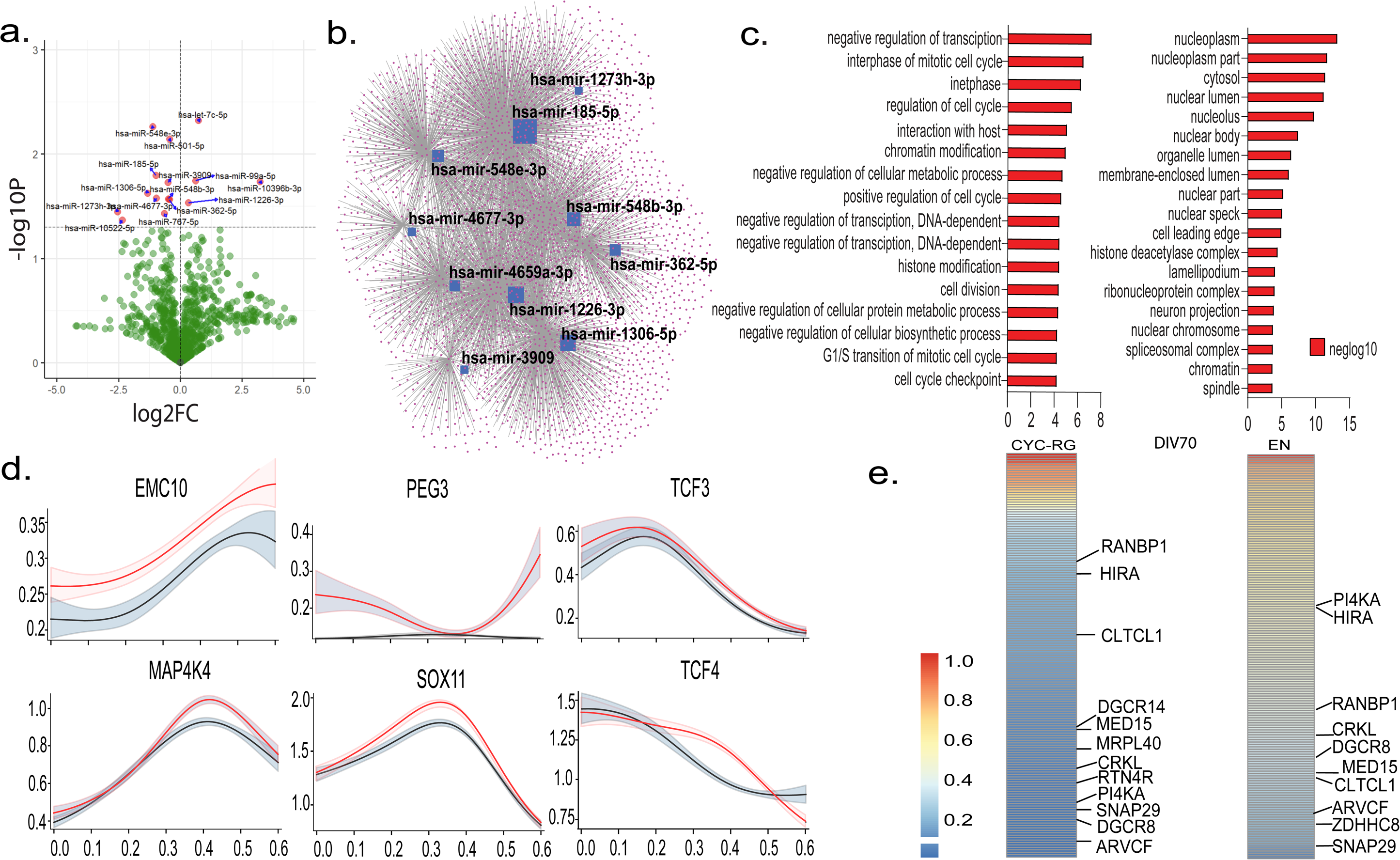
22q11.2-associated genes and genetic pathways underlying altered developmental pace. **(a)** Volcano plot depicting differentially expressed miRNAs in whole patient organoids at DIV70. **(b)** mirNet analysis of the network of downregulated miRNAs in patient organoids **(c)** GO term enrichment of mirNet-T computed targets of downregulated miRNAs. Targets are enriched for nuclear genes associated with cell cycle and cell division. **(d)** Latent time analysis of a subset of putative target genes of downregulated miRNAs using scVelo and CellRank; y axis denotes expression level; x axis denotes latent time. **(e)** PPI network analysis of dysregulated genes in patient RG-div (*left*) and ENs (*right*) with scoring of the contributions of the 22q11.2 locus genes to the network.

Reduction in miRNA expression levels is largely associated with up-regulation of target genes. Consistently, in RG-div, the major cycling neural progenitor population in our dataset, we found that predicted target genes of downregulated miRNAs have a significant overlap with the cell cycle and proliferation genes that are upregulated in RG-div of patient organoids (24% of upregulated genes are putative miRNA targets, **Supplemental Table 8**). No common cellular/molecular processes are enriched among predicted miRNA target genes downregulated in RG-div. Functional enrichment analysis of miRNA targets overlapping with upregulated DEGs along the entire EN lineage trajectory identified additional GO terms in early RG (RG-early) (“RNA metabolism” and “gene expression”), intermediate progenitors (“RNA metabolism” and “neurogenesis”) and EN (“RNA metabolism” “gene expression” and “neurogenesis). Interestingly, many of the upregulated EN DEGs that are predicted targets of the affected miRNAs are identified as negative regulators of neurogenesis or are associated with more immature neurons (such as *EPHA4, KDM1A, KIF5B, MAP4K4, NTRK3, PAX6, SLITRK5, SOX11, SOX4, TCF4, VIM*). No significant GO terms enrichment was observed among downregulated DEGs that are predicted targets of affected miRNAs in all these cellular populations (**Supplemental Table 8**).

We also tested the overlap between miRNA target genes and those in the DIV70 neural progenitor WGNA modules upregulated in patient EN lineage (**Supplemental Table 8**). We found that 20% of module 2 and 26.5% of module 9 genes are predicted targets of the downregulated miRNAs. This overlap was significant for module 9, which is associated with regulation of proliferation (*P* = 0.03, 1000 permutations). These results suggest that miRNA dysregulation may disrupt genes that maintain progenitors in a cycling state and hinder their differentiation.

We evaluated disparities in temporal trajectories of miRNA targets between patient and control organoids. Trajectory differences due to compromised miRNA control can be generated by genuine increase or failure to decrease transcript abundance. We looked at the expression trajectories of DEGs in EN lineage cell types that were also identified as target genes of downregulated miRNAs along the latent time. We identified 3 main patterns of trajectory divergence between patients and controls **(Fig 6d**). First, genes that are upregulated in patient progenitors and remain upregulated over time as exemplified by *EMC10* (a regulator of membrane protein trafficking whose upregulation impedes the acquisition of full neuronal functionality in patient neurons and animal models^22,38,39^) and *PEG3* (a cell cycle regulator that is typically highly expressed in progenitor cells but not in differentiated neurons^40^). Second, genes that show upregulation in progenitors but the changes dissipate over time as exemplified by *TCF3* (a WNT signaling antagonist that increases NPC self-renewal and represses neuronal differentiation^41,42^). Third, genes that show an upregulation at intermediate or late developmental timepoints but not at early progenitors as exemplified by *MAP4K4, SOX11* and *TCF4* (genes shown to inhibit neurite growth in differentiated neurons^43,44^).

Finally, we estimated in an unbiased fashion, the contribution of additional 22q11.2 genes to the signaling pathways and networks underlying the affected biological processes. We first derived a Protein-Protein Interaction (PPI) network of all DEGs (see Methods) for each affected cell type in the EN lineage using the STRING database^45^. We conducted a centrality analysis of each PPI network to rank the relative importance of each node using centrality scores (see Methods) and determined the 22q11.2 genes that are important for the network. For the PPI network in RG-div, we found that RANBP1^46^ and HIRA^47^ had higher scores. For the PPI network in EN, we found PI4KA ^48^and HIRA as the highest-scoring genes (**Fig 6e**). Despite limitations of this approach (i.e proteins such as DGCR8 that exert their effects via transcriptional and translation regulation instead via interacting with other proteins may not be weighted equally in the network), these results suggest that alterations in discrete sets of 22q11.2 genes are involved in deviations of protein interaction networks that contribute to distinct cellular and temporal aspects of neurodevelopmental trajectories. Overall, our analysis supports the notion that the high risk of atypical early neuronal maturation programs in patients is the result of concerted (parallel and sequential) actions of multiple genes on focal mechanisms instead of the effects of a single gene or pathway deficit.

### Aberrant gene expression in other cell types

While ENs are often the primary focus in studies of 22q11.2 deletion syndrome, there is growing recognition of the critical role of other cell types in this pathology^49–52^. Notably, our analysis reveals abnormal developmental trajectories and gene expression patterns in the IN lineage (**Supplemental Tables 4, 5**), indicating complex neurodevelopmental disruptions. Specifically, genes dysregulated in patient DIV150 INs are primarily involved in “neuron projection morphogenesis”, “regulation of transcription”, “regulation of cell cycle” as well as “regulation of glycolytic processes” indicating potential disruptions in neuronal development, cellular homeostasis, and metabolic pathways critical for interneuron maturation and function. The presence of INs in dorsal brain organoids, while previously described^49,51,52^, raises questions about their origins^53^. Recent evidence suggests an additional locus of interneuron generation in the human dorsal cortex^54^, but the majority of cortical interneurons are typically generated in ventral regions^55,56^. Our findings indicate that 22q11.2 hemizygosity results in additional defects in cortical interneuron development and potentially their function, underscoring the importance of ventral brain organoid models to further investigate interneuron dysfunction in 22q11.2DS^57^.

Additionally, we observed significant gene expression alterations, but no changes in cell abundance, in non-neuronal cell types such as astrocytes and the choroid plexus (**Supplemental Tables 2-5**), indicating that the 22q11.2 deletion affects developmental processes beyond neuronal populations. In patient DIV150 astrocytes, for example, dysregulated genes are primarily involved in ‘neurogenesis’ and ‘mitochondrial ATP synthesis coupled electron transport.’ These findings suggest potential interactions between neurons, glial cells, and cerebrospinal fluid (CSF), highlighting the need for further investigation into the broader implications of these deletions on brain development and function.

### Aberrant gene expression in EN lineage displays signatures of neuropsychiatric genetic liability

The growing appreciation that childhood neurodevelopmental disorders and the early developmental antecedents of SCZ may share genetic liability^58^ prompted us to ask whether the aberrant transcription program underlying impaired maturation contains relevant genetic signatures. Specifically, we asked if the aberrant transcripts in the EN lineage of patient organoids involve genes identified as harboring rare coding damaging variants in exome sequencing analysis of patients with SCZ, autism spectrum disorders (ASD) and developmental delay/intellectual dysfunction (DD-ID). To achieve adequate statistical power, we analyzed DEGs in cell clusters with at least 100 DEGs at both DIV70 and 150 using the DNENRICH pipeline and curated reference patient datasets (see Methods) for genes carrying rare LoF variants, including genes very likely to be intolerant to LoF variations as measured by the probability of LoF intolerance (pLI) score^59^, which ranks genes from most mutation tolerant (pLI = 0) to most intolerant (pLI = 1) and is widely used to prioritize candidate disease genes. Genes carrying rare synonymous variants were used as a control.

Our rare-variant burden analyses revealed a significant overlap between the expression changes in genes harboring LoF variants and a wide range of cell types in the EN lineage, particularly at later developmental stages. (**Fig 7**). Notably, DEGs harboring LoF variants in SCZ cases are most consistently enriched in EN (including genes such as *GRIA3, BOD1L1, EIF2S3, FABP7, POTEF, SLITRK5, ZMYM2, ANKRD12,* KDM6B) whereas DEGs harboring LoF mutations in ASD are most consistently enriched in neural progenitors (IPC-nEN) (**Fig 7a)**. For all three disorders, the set of highly mutation intolerant (pLI≥0.9) DEGs harboring rare LoF variants displays a wider cellular representation within the EN lineage. (**Fig 7b)**. As expected, enrichment signals are absent for synonymous variants (**Fig 7c)**. Our findings suggest that during early brain development transcripts dysregulated in differentiating neurons of 22q11.2 deletion carriers are enriched for products of genes that when mutated increase the risk for SCZ and neurodevelopmental disorders.

**Fig 7:**
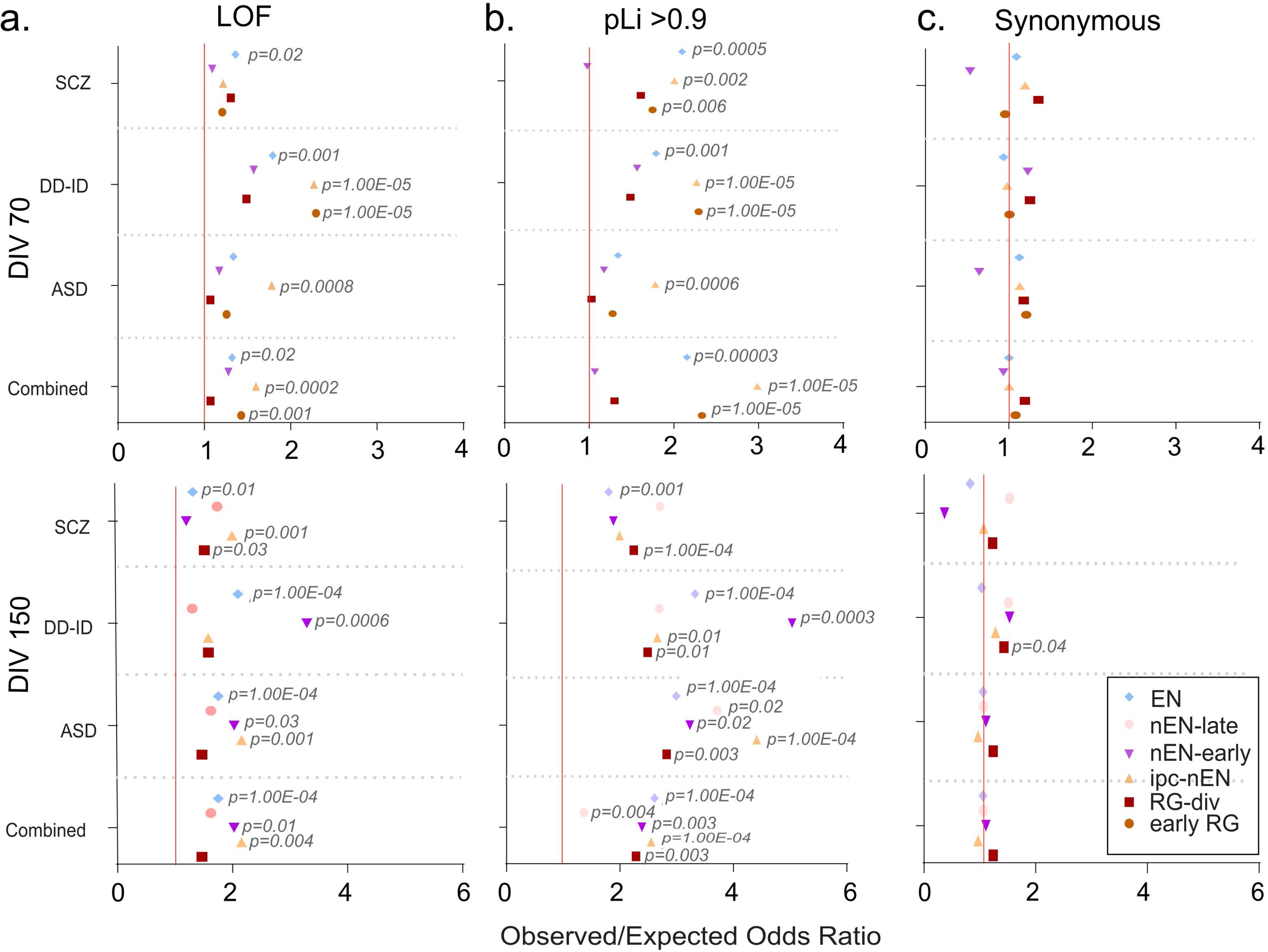
Overlap between genes harboring rare LoF risk variants and EN lineage cell types. Results from permutation-based analysis of overlap between EN lineage cell types and DEGs harboring rare coding damaging variants at DIV70 (*top*) and DIV150 (*bottom*) using the DNENRICH pipeline and curated reference datasets. Overlap between cell types in the EN lineage and dysregulated genes harboring LoF risk variants **(a)**, highly intolerant LoF variants (pLI ≥ 0.9) **(b),** or synonymous variants **(c).**

## DISCUSSION

In this study, we generated forebrain organoids from iPSCs derived from patients with 22q11.2DS and SCZ. To minimize variability due to the heterogeneity in disease penetrance and clinical presentation, we selected iPSC lines from patients with the same 3Mb deletion and similar clinical severity. Genetically related, age-matched sibling pairs were also included for comparison. It has been shown that key variability in iPSC-based models arises from differences across individuals^60–62^. To account for this, we prioritized the analysis of multiple patient iPSC lines, thereby improving the generalizability of disease phenotypes while controlling for genetic background variability. Our PCA analysis confirmed that genetic ancestry and genotype are the primary contributors to transcriptional variance in cortical neurons, validating that our experimental design effectively captures phenotype variability while minimizing confounding from hiPSC line variability.

In the context of 22q11.2DS, previous studies have begun to address the neurodevelopmental consequences of the 22q11.2 3Mb deletion in patient-derived neuronal cultures^23,63–67^. However, our study is the first to detail the developmental trajectory of iPSC-derived neurons from patients with the 22q11.2 3Mb deletion and SCZ under biologically relevant conditions. By leveraging a forebrain organoid culture system that supports spontaneous differentiation into diverse cell types, we provide a more representative model of neurodevelopmental processes. The ability of patient and control organoids to faithfully recapitulate the expected 22q11.2 gene expression profile, combined with consistent molecular and cellular findings across organoid batches, rules out random developmental variability as the source of the observed aberrations.

Single-cell RNA sequencing (scRNA-seq) and orthogonal experimental validation revealed a delay in neuronal differentiation and maturation in patient organoids during periods of active neurogenesis, DIV70 and DIV150 of organoid development, which strongly correspond to the fetal human telencephalon between 7-37 weeks post-conception^53^. Specifically, scRNA-seq quantification of EN lineage cells showed a nominally significantly reduced proportion of differentiated neurons in patient-derived organoids compared to controls. Pseudotime trajectory analysis of DIV70 and DIV150 EN lineage revealed a diminished representation of patient cells in the terminal, mature neuron phase of the trajectory. In agreement with the scRNA-seq data, we observed a marked reduction in the fraction of mature (CTIP2+) neurons in patient-derived DIV20 neurospheres and a significant decrease in cortical plate (CP) thickness in patient DIV45 organoids compared to controls. Bulk RNA-seq on sorted neurons identified a significant enrichment of downregulated synaptic genes, with no such enrichment among upregulated genes, indicating a delay in synapse maturation. Consistent with this, gene expression profiling by scRNA-seq revealed that genes downregulated in patient-derived DIV70 EN cells were primarily involved in neuronal differentiation and maturation, while upregulated genes were enriched for pathways related to the negative regulation of neurogenesis, neuron growth, and axon guidance. These molecular disturbances were paralleled by morphological abnormalities, such as sparser neurites and electrophysiological deficits, including reduced amplitude of glutamate-induced calcium transients.

In line with the marked decrease in mature neuron production, our 3D immunostaining analysis of DIV70 organoids revealed a significant increase in the proportion of actively proliferating NPCs (Ki67-positive/SOX2-positive) in patient-derived organoids, consistently observed across all control and patient pairs. However, the total number of NPCs (SOX2-positive) remained unchanged. While scRNA-seq quantification of EN lineage cell types showed a nominally significant higher number of dividing progenitors, pseudotime trajectory analysis as well as calculation of velocity streams and differentiation potential in the DIV70 datasets indicated that EN lineage cells in patient organoids tend to remain in progenitor states longer compared to controls. Moreover, our neurosphere and PIP-FUCCI sensor assays confirmed that patient NPCs exhibit a lower proliferation rate and slower differentiation into neurons compared to controls, as evidenced by a lower proportion of cycling NPCs in the S phase, an accumulation in the G1 phase, and a reduced number of neurons in patient samples. Proliferation deficits may be masked by delayed cell cycle exit in patient NPCs at later developmental stages, when neurogenesis becomes predominant, ultimately resulting in a total patient NPC count comparable to controls. Further supporting our conclusion that patient NPCs exhibit altered cell cycle dynamics, scRNA-seq-based gene expression profiling and WGCNA revealed that genes upregulated in patient DIV70 RG-div, the primary dividing NPC population, were predominantly involved in cell cycle regulation and the promotion of cell division. In contrast, downregulated genes were associated with neurogenesis and neuronal differentiation. Together, these orthogonal results suggest delayed differentiation of progenitors in patient organoids. The minor discrepancies between immunostaining and scRNA-seq-derived findings may reflect broader challenges in reconciling scRNA-seq-derived molecular signatures, which distinguish cell types based on co-expression of rare or low-expression transcripts, with classical immunostaining, which relies on more abundant, well-characterized protein markers. This distinction is particularly important when examining dynamic transitions between cellular states (neurogenesis) versus stable endpoint states (mature neurons).

Interestingly, analysis of cell distribution along the pseudotime trajectory also suggests that the 22q11.2 deletion may delay acquisition of neuronal fate as early as the radial glial stage, prior to the onset of neurogenesis. This interpretation aligns with recent findings that the rate of neuronal maturation in humans is predetermined before neurogenesis, through the establishment of a ‘barrier’ in progenitor cells formed by various chromatin regulators, which is then propagated to newborn neurons^68^. However, additional disruptions caused by the 22q11.2 deletion along the temporal progression from neural precursors to post-mitotic neurons may further impede the acquisition of full neuronal morphology and functionality^18^.

The timing, magnitude, and spatial distribution of delayed neuronal maturation likely leads to regional changes in brain structure, connectivity, and activity patterns early in brain development. These changes may propagate to later prenatal and postnatal stages, contributing to the behavioral and cognitive symptoms associated with 22q11.2 deletions. Brain imaging studies in children with 22q11.2 deletions report region-specific alterations in cortical thickness and surface area across several brain regions^69–73^. During brain development, cortical thickness and surface area are affected by complex interactions among progenitor cell divisions, neuron growth and neurite branching^74–78^. Therefore, it is plausible that the altered proliferation dynamics and expansion of progenitor cells, coupled with impaired neuronal growth and differentiation (as observed in patient organoids), could contribute to the reported imaging findings. However, the precise mechanisms connecting these neurodevelopmental alterations with the observed brain imaging differences remain to be clarified.

Transcriptomic analyses of key cell types highlight that 22q11.2 deletions lead to pervasive and improperly orchestrated gene expression alterations in a wide range of cell types and ensembles of genes across the entire EN lineage. This analysis underscored biological pathways crucial for neuronal development and function that are affected by 22q11.2 deletions during human early brain development. Previous studies using mouse models of the 22q11.2 deletion have established a clear link between altered miRNA biogenesis and the resulting behavioral and circuit changes associated with this genomic alteration ^20,36,38^. Specifically, Dgcr8 haploinsufficiency has been shown to significantly contribute to the transcriptional changes in the adult brain of these mutant mice ^20^. Furthermore, the 22q11.2 deletion leads to abnormal miRNA processing in monolayer cultures of patient iPSC-derived neurons, with functional consequences on their transcriptome as well as electrophysiological and synaptic properties^23,63,79^. In alignment with these findings, our analysis of patient-derived organoids revealed that miRNA dysregulation can affect gene expression in a cell-type specific manner, altering both the levels and the temporal dynamics of genes critical for neuronal development and maturation. These disruptions in gene expression patterns may hinder the initiation and progression of maturation programs, echoing existing evidence that miRNAs play crucial roles in regulating temporal transitions during development^80–82^, including in human brain development^83^. This underscores the importance of understanding miRNA dynamics in the context of 22q11.2 deletions, as they may be pivotal in mediating the neurodevelopmental impacts of this genomic lesion. Noteworthy targets of miRNA dysregulation include epigenetic regulators such as EZH2, a key component of a recently identified epigenetic barrier to maturation, which is established before the onset of neurogenesis^68^. EZH2 was upregulated in RG-early of DIV70 patient organoids (*Padj* = 0.001, **Supplemental Table S2**). This raises the interesting possibility that the effects of miRNA dysregulation at this early stage may be mediated, at least in part, via the epigenetic machinery. Another noteworthy target is the EMC10 gene, which shows an altered temporal expression trajectory across the entire EN lineage at DIV70, with significant upregulation in patient RG (*P* = 1.56×10^-6^), IPC-nEN (*P* = 5.3×10^-5^) and EN (*P* = 7.3×10^-6^) but not in astrocytes, choroid or interneuron lineage cells (**Supplemental Table 2**). Indeed, effective knockdown of EMC10 to near wild-type levels in patient iPSC-derived neurons restores key morphological and functional EN alterations caused by 22q11.2 deletions^22^. Furthermore, normalization of Emc10 levels in either the developing or adult mouse brain rescues several cognitive deficits associated with 22q11.2DS^22,31^. These findings establish a clear link between miRNA regulatory mechanisms and aspects of the clinical manifestations observed in 22q11.2 deletion carriers and they highlight the potential of transcriptomic changes in patient organoids to serve as a pool of therapeutic targets with translational value.

While altered miRNA biogenesis may affect the pace of neuronal development, it is likely not the sole driving factor. Findings from both human genetic and animal model studies suggest that the deficits observed in 22q11.2 deletion carriers result from the cumulative and concerted actions of multiple genes within the deleted region, acting over developmental time and space^8,84,85,86^. This is supported by our PPI (protein-protein interaction) centrality analysis, which indicates that 22q11.2 genes, in addition to DGCR8, may make complementary contributions to the altered signaling pathways in distinct cell types and developmental stages. We identified *RANBP1, HIRA* and *CLTCL1* as key candidates underlying perturbations of protein networks at the early radial glial stage. Similarly, HIRA and PI4KA were identified as key component of regulatory protein networks affected in embryonic EN. These findings suggest that a multi-level experimental approach is needed to understand the integrated contributions of the deleted genes, which could guide therapeutic interventions.

Our results are in good concordance with prior studies of iPSC-derived neurons from patients with the common 22q11.2 3Mb deletion, particularly with respect to disrupted EN maturation^23,63–67^. However, some discrepancies were observed, likely reflecting variations in experimental conditions, differentiation protocols, or patient-specific factors, which warrant further investigation. Specifically, proliferation assays of iPSC-derived NPCs obtained from controls and patients with 22q11.2 deletion syndrome diagnosed with a psychotic disorder showed a trend toward decreased proliferation^64^. Additionally, previous analyses of neurospheres derived from hiPSCs of two patients with the 22q11.2 deletion and SCZ also provided evidence of impaired neurogenesis and neurite outgrowth^65^. While not explicitly analyzing neurogenesis, Nehme et al.^67^ performed RNA sequencing during three stages of EN differentiation using iPSCs from 22q11.2 deletion carriers (many with SCZ) and controls, employing an NGN2 overexpression protocol to induce neuronal differentiation. They identified significant alterations in genes and pathways related to proliferation in NPCs as well as to activity-dependent gene expression and synaptic biology (in mature neurons), consistent with impaired synaptic maturation. Functional assays confirmed reduced synaptic functionality in 22q11.2 deletion neurons. In contrast, Li et al. ^66^using a similar 2D neuronal differentiation protocol, reported no synaptic deficits but instead highlighted mitochondrial impairments, including reduced mitochondrial DNA-encoded proteins, ATP production, and complexes I and IV activity. In our dataset, dysregulation of genes encoding mitochondrial membrane proteins was primarily observed in progenitor neurons (nENs) (**Supplemental Table 3**).

A recent study^23^ using scRNA-seq analysis of cortical spheroids derived from two patients with the 22q11.2 deletion (SCZ diagnosis unknown) and two controls reported similar cell diversity across groups, suggesting no major defects in corticogenesis. The reasons for the discrepancy with our findings are unclear but likely multifactorial. Potential contributing factors include differences in clinical phenotype severity, the small sample size, and suboptimal matching of patient and control lines by age, sex, or genetic background, which may obscure transient developmental alterations. Additionally, differences in experimental conditions and developmental stages between the two studies may have influenced the results. Indeed, re-analyzing the scRNA-seq data from Khan et al.^23^ using our pipeline and, after standard QC, we found very few radial glia-dividing (RG-div) cells—suggesting their cortical spheroids were at a more mature stage, with diminished proliferative features. Comparing our bulk RNA-seq DEGs with theirs revealed minimal overlap outside the deletion region (7 genes: *PALM, SLC39A12, LRWD1, ZNF502, MMP17, AC093866.1, AC020604.1*), likely reflecting differences in cell composition, as their data came from whole organoids and ours from purified cortical neurons. Among shared DEGs, we identified *PALM*, involved in palmitoylation-dependent synapse formation^87^, which is particularly relevant given the deletion of *ZDHHC8*, a palmitoyl transferase, in 22q11.2DS^19,88^. Comparing their top 50 excitatory neuron DEGs to our 442 revealed 8 overlapping genes (*CNTNAP2, NTS, GPM6A, GRIA2, SOX4, NSG1, NTM, LY6H*), many involved in synapse and neuronal projection function. Notably, Khan et al.^23^ also reported blunted calcium signaling in patient-derived cortical neurons, consistent with our findings.

In the context of SCZ, the primary psychiatric manifestation associated with 22q11.2 deletions, our findings align with reports of aberrant NPC proliferation and impaired differentiation observed in SCZ patient iPSC-derived neuronal and neural organoid models (reviewed in ref^59^). However, it is important to note that many of these studies face challenges in addressing confounding factors, such as genetic heterogeneity, and, in some cases, in clearly identifying the precise identity and type of neurons being assessed. Existing findings suggest that NPCs struggle to properly differentiate into mature neurons, indicating a mechanism where either a smaller initial NPC pool or impaired differentiation contributes to reduced neural density later in the disease, with the underlying molecular causes varying among patients due to the genetic heterogeneity of the condition^89,90^. Additionally, differentiated glutamatergic ENs exhibit defects in neurogenesis and morphological changes, such as impaired dendritic arborization, which are further associated with functional impairments, as patient neuronal subtypes frequently demonstrate reduced synaptic connectivity^91,92^. Interestingly, these results contrast sharply with recent studies showing accelerated maturation of the same neuronal lineages in brain organoid models carrying mutations in specific autism spectrum disorder (ASD) risk genes^93^. This divergent pattern of maturation is in line with human genetic studies that have repeatedly identified 22q11.2 deletions as top hits in studies of SCZ but not ASD genetic risk^94^.

Importantly, previous studies have shown that SCZ risk genes targeted by rare *de novo* mutations often exhibit highest expression during the prenatal period^95^. Many prenatally-biased risk genes are highly expressed during the first and second trimester of pregnancy, a period of active cortical neurogenesis and a vulnerable time for brain development due to highly dynamic changes in gene expression, high level expression of genes intolerant to mutations and increased susceptibility to environmental insults^83,96,97^. Consistently, we have shown that genes carrying SCZ rare risk variants are enriched among DEGs in patient organoid neurons. This supports the notion that, for a fraction of patients with SCZ, genetic liability may affect the pace of early brain developmental trajectories similar to the ones described here. In that context, our findings may be generalized to other genetic causes of SCZ, especially ones with large effect mutations and have the potential to facilitate the pursuit of preventive and therapeutic interventions.

## METHODS

### hiPSC generation and characterization

Q5, Q6, Q1, and Q2 iPSC lines were generated as described previously^22,98^ at the Columbia Stem Cell Core via non-integrating Sendai virus-based reprogramming of monocytes from a donor with 22q11.2DS and SCZ and a healthy sibling control. We confirmed that all iPSC lines maintain their stemness using RT-PCR and immunostaining for stem cell markers (NANOG, OCT4/SOX4, SSEA-4 and TRA-1–60^22^. Given the importance of genetic testing of iPSC lines^99,100^, we confirmed their genomic stability through G-banding karyotype analysis and verified the genotypes and size of the 22q11.2 deletion using a multiplex ligation-dependent probe amplification (MLPA) assay^98^. Clinical information on patients is provided in ref^101^. Q1/Q2 is pair of siblings; Q5/Q6 is pair of dizygotic twins discordant for the 22q11.2DS and SCZ. The newly generated iPSC lines have been deposited in the NIMH repository. QR19, QR23, QR20 and QR27 lines were obtained from the NIMH Repository and Genomics Resource (http://www.nimhstemcells.org/)^64^. Usage of these iPSC lines has been approved by the Columbia University Embryo and Embryonic Stem Cell Research Committee (ESCRO). Sample usage in downstream analysis is summarized in **Supplemental Table 10**.

### Generation of dorsal forebrain cerebral organoids

We generated dorsal forebrain patterned organoids using a protocol that was adapted from Kadoshima et al^21^. At DIV 40, organoids were transferred to maintenance media containing 1% N2 and 2% FBS. After DIV 80, they were maintained in media containing 1% N2, 2% FBS and 1% B27 supplement.

### Bulk RNA-seq

We used a set of patient and control organoids at DIV70 (8-10 organoids per line, 4 case and 4 control lines). The organoids were transduced with the AAV-hSYN-GFP virus two weeks before DIV70 (Addgene, 105539-AAV1). We dissociated the transduced organoids using papain digestion (Worthington, LK003160) and sorted the GFP positive neurons by FACS. Neurons were lysed in TRIzol reagent (Thermo Fisher Scientific, 15596026) and total RNA was extracted from the cells using Qiagen RNeasy Kit (Qiagen, 74104). Quality control checks using a Bioanalyzer (Agilent 2100 bioanalyzer) were performed to ensure the RNA Integrity Number (RIN)>9.0 for all samples. Library preparation and RNA sequencing were conducted to generate 20 million paired-end reads at the Columbia Genome Center on Illumina Novaseq 6000 instrument using STRYPOLYA library prep kit. Raw reads were processed and aligned to human genome (hg38) using the STAR alignment tool^102^and the gene count matrix generated was analyzed using the DeSEQ2 pipeline to identify the DEGs^103^. The genes with LFC>0.2 and an adjusted P<0.05 were considered as DEGs in downstream analysis. PCA analysis was conducted using iDEP^104^. GO term enrichment was performed using the G:Profiler tool^105^. The PPI network was analyzed using string-db (https://string-db.org).

### miRNA-seq

Total RNA extraction with miRNA enrichment was performed using the Vana™ miRNA Isolation Kit (Thermo Fisher Scientific, AM1561). Total RNA quality and quantity was assayed with Bioanalyzer 2100 (Agilent, CA, USA), with RIN number >7.0. Approximately 1 Dg of total RNA was used to prepare small RNA library according to the protocol of TruSeq Small RNA Sample Prep Kits (Illumina, San Diego, CA, USA) and single-end sequencing 50bp on an Illumina Hiseq 2500 (following the vendor’s recommended protocol) was conducted as described in ref^101^. For the GO-term enrichment analysis of the target genes of differentially expressed miRNAs (153/133, DEmiRs), the webtool miRNet 2.0 (https://www.mirnet.ca/) with standard settings (tissue: all, targets: genes [TargetScan]) was used^106^. Analysis of the overlap of dysregulated genes and miRNA targets was performed by converting the gene lists to their corresponding genomic coordinates. To assess the significance of the overlap between gene sets, we extracted the genomic coordinates of all genes that were expressed in the organoids (determined by their expression levels in scRNA-seq) as the search space and used the randomization function in RegioneR to “resample” the regions that only contain the genes in the search space^107^. One randomly “resampled” region set had the same size as the dysregulated genes, while the other randomly “resampled” region set had the same size as miRNA targets. The number of overlaps between these two region sets was determined by RegioneR and 1000 permutations were performed to generate a frequency distribution of random overlaps between two region sets to determine the empirical p value of the actual overlaps.

### scRNA-seq

We generated a high viability (>90%) single cell suspension from 3 pairs of 70-day and 150-day old organoids, a total of 6 distinct lines with 3 organoids per line at two time points. The single cell suspension was sequenced on the 10x Genomics platform at the Columbia Genome Center. Our analysis workflow included (i) aggregation of all data (12 datasets from 6 samples and two time points) with depth normalization^108^ (ii) Quality Control (Scanpy) ^24,109^ (iii) Normalization (Scran^110^), (iv) Integration (LIGER^111^), (v)Clustering (Leiden^25^), (vi) Differential Gene Expression Analysis (vii) Visualization (UMAP^112^) (viii) Cell type annotation (SingleR^27^). We used a recently published dataset that classified the signature transcriptomic profiles of the cell types within the human fetal forebrain at various stages of development^26^, as a cell type signature reference to annotate the cells in our data. We used a custom-built stratified subsampling approach to validate cell type annotations and calculated the percentage of times a particular annotation is assigned to a cell over multiple random training permutations on stratified subsamples (https://github.com/BinXBioLab/scRNAseq). Next, we defined high-confidence annotations as those that had a higher percentage than random assignments and ensured that 98% of our cell type annotations were high-confidence. The number of cells in some neuronal subtypes is too small, limiting the scope for further analysis. To address this, we grouped such subtypes into their major cell types to increase statistical power. Additionally, we excluded cell types with a total cell count of fewer than 50 from further analysis. We compared cell type abundance differences using Milo^28^. We conducted an analysis of DEGs in various cell type populations using a significance testing under the Hurdle model implemented the MAST pipeline^33^ to calculate the p value and Coeff (log odds of expression) between genotypes..

For WGCNA analysis of control organoids at DIV70, we converted the scRNA-seq dataset into Seurat format, subset cells in the EN lineage and used the scWGCNA pipeline with default settings and using top 2000 variable genes for the analysis^35^. After the WGCNA modules were identified, we scored the expression of module genes in both patient and control datasets at DIV70 using the “scanpy.tl.score.genes” function and compared the expression scores of these WGCNA modules using a linear model with each individual iPSC line as random variable and genotype as a predictor as described previously^93^.

To assess developmental trajectories, we employed two approaches. First, we used Monocle3 ^113^ to assess pseudotime trajectories of different developmental lineages in our scRNA-seq dataset. For each lineage and DIV, we plotted Ridgeline plots of cell distribution along pseudotime and performed the KS test to assess if the distributions were significantly different between patient and control organoids^93^. Second, we used the scVelo tool ^31^ to analyze RNA velocity of individual cells in batch-corrected patient and control organoid datasets as a whole, followed by an analysis of differentiation potential and absorption probability using the CellRank tool^30^. Latent time represents an internal clock of each cell based on its transcriptional dynamics and reveals the time a cell stays in the current status before it transits to the next cell fate status.

For analysis of transcription factor regulatory networks, we used the pYSCENIC pipeline^114^. First, we generated a representation of the DIV70 dataset in TF space and confirmed that our cell representation is consistent with cell-type annotation within this space. For example, we could detect enhanced activity of the neuronal transcription factors NEUROD1 in the neuronal clusters while the stem and progenitor cell marker SOX2 has a higher activity in the radial glia and dividing progenitor clusters. After we identified differentially active TF between patients and controls in various cell types, we compiled the top differentially active TF and their downstream effector genes (regulons) and conducted GO term enrichment analysis on the effector genes. GO term enrichment of DEGs obtained from the above analysis was performed using G:Profiler^105^

### Analysis of cell proliferation features

First, to assess the proportion of dividing cells, organoids were fixed as described above. Organoids were subjected to 3D immunostaining using a protocol modified from the iDISCO whole-mount staining protocol (https://idisco.info/idisco-protocol/) that allowed 3D imaging of a large area and depth (200µm) of the organoid. Samples were washed, permeabilized in 0.2% TritonX-100 for 24 hours at 37°C and blocked in 10% Horse serum and 0.2% TritonX-100 for 24h at 37°C. Primary antibodies were then added along with 10ug/ml Heparin, 3% Horse serum and 0.2% Tween-20 for 50 hours at 37°C. Primary antibodies used for cell cycle analyses were rabbit anti-Ki67 (Abcam, ab15580, 1:200) and goat anti-SOX2 (R&D Systems, AF2018-SP, 1:200). Samples were then washed and treated with secondary antibodies (donkey secondary antibodies Jackson Immunoresearch) at 1:300 dilution for 36 hours. Samples were then incubated overnight with DAPI, washed and then mounted in RapiClear medium (SunJin Labs, RC149002) in 18 well chamber slides (Ibidi, 8187) overnight for clearing and imaging on W1-Yokogawa Spinning Disk Confocal (Nikon Instruments, Tokyo, Japan) at 40x magnification. To analyze the images, the DAPI channel was first segmented using ilastik^115^ to provide a binary segmentation mask of the nucleus foreground from the background. Binary erosion of the DAPI mask was performed on Fiji^116^ to remove noise points. The “distance transform watershed 3D” method from MorphoLibJ^117^ on Fiji was used to extract and distinguish individual DAPI nuclei and assign them unique mask ID. Within the cell mask of each individual DAPI nucleus, the signal in other channels (SOX2, Ki67) was measured using customized Python code and a cut-off threshold was applied to determine cell identity. The thresholds were manually chosen based on each image and were further proofread. Cell proportions were compared using the KS test, p<0.05 was the significance threshold. Second, to determine the proportion of cycling cells in each of G1, S and G2/M phase, we employed a lentiviral PIP-FUCCI biosensor^118^ from Addgene (pLenti-PGK-Neo-PIP-FUCCI plasmid #118616). Concentrated viral supernatant was transduced into 2D culture purified NPCs overnight. Fresh medium was then replaced and NPCs were further cultured for 5 days before fixation with 4% PFA. Imaging was performed using an Olympus iX81 inverted microscope using 10x objective lenses with DAPI, Venus and mCherry channels. Cell segmentation and counting were performed using a modified protocol with the TrackMate plugin in Fiji^119^. The difference between the percentage of patient and control NPCs at each cycling phase was determined by a student t-test. All experiments were repeated three times. Third, to determine the neurogenesis features of NPCs, a FACS-based analysis was conducted. Specially, DIV 20 NPCs were cultured in the differential medium for 10 days and then first dissociated into single cell suspensions using papain digestion (Worthington Biochemical), followed by fixation and permeabilization with eBioscience™ Foxp3/Transcription Factor Staining Buffer Set (Thermofisher). Primary antibody (SOX2, CTIP2) incubation was conducted at 4 °C overnight. Following a washing step with PBS, cells were incubated with 1:500 Alexa Fluor secondary antibodies (Thermofisher) for 1hr at room temperature. Analysis was performed on a NovoCyte (Penteon) flow cytometer. 100-500K events were acquired for each sample with fluorescence measured in logarithmic scale. Cells incubated only with secondary antibodies served as background fluorescence, and were used to set the gating parameters. After exclusion of cell aggregates and small debris by forward and side scatter gating, data were analyzed using the NovoCyte software and plotted in a histogram format. Fluorescence gates were set below 2% of blank histogram and events corresponding to a fluorescence signal exceeding this percentage were considered as positive events. The difference between percentages of patient and control NPCs at each cycling phase was determined by a two-sided student t-test.

### Analysis of neuronal morphology

We analyzed organoids at DIV70. At DIV55, we infected organoids for 2 weeks with the AAV1-hSYN-eGFP virus (Addgene, 105539-AAV1), which directs expression of eGFP under the neuron-specific synapsin promoter. At DIV67, we placed the organoids onto a glass coverslip and immobilized them in a droplet of 50% Matrigel in culture media (Corning, 354277). At DIV70, organoids were fixed in 4%PFA (Biotium, 22023) for 3h at RT and then blocked for 1.5h at RT in 0.3% Triton-X and 10% horse serum (catalog#H0146, Sigma-Aldrich, St. Louis, MO, USA) solution. After fixation, organoids were stained with the primary got anti-GFP antibody (Rockland, 600-101-215, 1:500) to amplify the fluorescence signal in a 0.1% Triton-X and 2% horse serum solution overnight at 4°C. Coverslips were then washed 3×15 min with DPBS (catalog#D8537, Sigma-Aldrich, St. Louis, MO, USA) and cells were incubated for 2h with the secondary antibody at room temperature followed by 3×15 min DPBS washing. Organoids were imaged as a whole mount on W1-Yokogawa Spinning Disk Confocal (Nikon Instruments, Tokyo, Japan) at 40x magnification to facilitate imaging of a large area of the organoid without sectioning and thus preserving the cell morphology. We generated 3D traces of individual neurons using Imaris (Bitplane). We quantified the number of branchpoints, number of primary neurites and total neurite length for each traced cell. We analyzed a total of 8-11 organoids per line over 3-4 separate batches generated on different dates. A total of 66-166 cells per iPSC line were analyzed at DIV70. In addition to immunostaining for GFP, in a subset of organoids, we performed immunostaining for the somatodendritic marker MAP2 (chicken anti-MAP2, Abcam, ab5392, 1:500) and immature neuron marker DCX (mouse anti-DCX, Santa Cruz Biotech sc-271390, 1:250) to assess cell development and neurite specification.

### Analysis of organoid growth

To analyze global organoid growth, organoids were imaged weekly using a bright field microscope (4x objective, Zeiss Primovert inverted microscope). Fiji^116^ was used to trace the border of the organoids and define regions of interest (ROIs). For each time point, 2 parameters were measured; the perimeter of the ROI and the area enclosed by the ROI. These two measurements were used to track organoid size over time.

### Ca^2+^ imaging

We constructed a novel Lentivirus vector driving GCAMP6s expression (Lenti-Ubi-mRuby2-GSG-p2A-GCAMP6s). Organoids were transduced with the Lentivirus vector 14 days prior to recording. Ca^2+^ activity as indicated by GCAMP6s fluorescence intensity was assessed visually prior to image acquisition. GCAMP6s signal was recorded for 25 min using an Olympus IX81 epifluorescence microscope at 4x magnification, with a frame rate of 1 frame per second (fps) and recorded for 1500 frames. At the 500 second mark, a glutamate solution was added to the culture medium at a final concentration of 100M. Ca^2+^ activity was measured in a set of organoids that had been cultured for days 90, 168, 238 and 259 to determine optimal timepoint for Ca^2+^ activities. Images were stabilized with a stabilizer plugin in ImageJ. Denoising, segmentation and Ca^2+^ event detection were performed using a CaImAn-based pipeline^120^. Ca^2+^ activities with DF/F local maxima value higher than five times the standard deviation from baseline were classified as one firing event. For Ca^2+^ event peak heights, only cells with temporally non-overlapping and distinguishable Ca^2+^ events were used, and only the Ca^2+^ event in the middle in temporal order was used in cases when one cell has multiple Ca^2+^ events.

### Analysis of the overlap between genes harboring rare LoF risk variants and EN lineage cell types

We used DNENRICH^121^ for calculating the permutation-based significance of gene set enrichment among genes hit by *de novo* mutations accounting for potential confounding factors such as gene sizes and local trinucleotide contexts. DNENRICH analyses were performed using 100,000 permutations. The following inputs were used for this analysis: 1) gene name alias file: the default dataset included in the DNENRICH package, 2) gene size matrix file: the default dataset included in the DNENRICH package (for RefSeq genes), 3) gene set file: the set of DEGs identified by the MAST analysis of individual cell types; all genes were equally weighted, 4) mutation list files: lists of genes (equally weighted) carrying LoF DNMs in SCZ, ID/DD, NDD and ASD from denovo-db SSC and non-SSC. For SCZ we also added significant and nominally significant DNMs from the SCHEMA dataset (p<0.05, https://schema.broadinstitute.org/results) and 5) background gene file: the default gene list in the gene size matrix. The numbers of LoF DNMs in SCZ, ID/DD, ASD were 1657, 1373, 997 and the respective numbers of synonymous mutations were 3610, 794, 1644. For analyses restricting the input LoF DNMs to those in genes intolerant to LoF mutations, we used pLI scores≥0.9^59^.

### Protein-protein interaction (PPI) network analysis

We used the search tool for retrieval of interacting genes (STRING) (https://string-db.org) database, which integrates both known and predicted PPIs, to seek potential interactions between DEGs and visualize functional interactions of these proteins. Active interaction sources, including text mining, experiments, databases, and co-expression as well as species limited to “Homo sapiens” and an interaction scoreD>D0.4 were applied to construct the PPI networks. A centrality analysis on each PPI network was conducted to estimate how important a node or edge is for the connectivity or the information flow of the network using CINNA (Central Informative Nodes in Network Analysis)^122^ R package.

### Cross-studies Comparison analysis

The RNA-seq DEG lists were obtained from the supplemental tables of the Khan et al^23^., study. DEGs were merged based on their gene symbols. Overlapping genes were identified, and GO term enrichment analysis was performed using ToppGene (https://toppgene.cchmc.org/enrichment.jsp) (**Supplemental Table 11**).

## Supporting information

Supplemental Table 1_DIV70_sorted_neuron_bulk_DEGs

Supplemental Table 2_DIV70_scRNAseq_DEGs

Supplemental Table 3_scRNAseq_DIV70_GO_enrichment

Supplemental Table 4_DIV150_scRNAseq_DEGs

Supplemental table 5_DIV150_scRNAseq_GO_enrichment

Supplemental Table 6_Tx regulons

Supplementary Table 7_scWGCNA modules

Supplementary Table 8_final_miRNAseq_DEGs

Supplementary Table9_Clinical and demographic characteristics

Supplemental Table 10 Sample Usage in Experiments

## Author Contributions

B.X., J.A.G. and S.B.R. contributed to the conception and design of the study; S.A.K, B.X. and J.A.G. supervised all the experiments; S.B.R., H.Z.,Y.S. and Z.S. contributed to characterization and maintenance of human iPSC line as well as generation and culture of organoids; R.J.S contributed to the collection and clinical diagnosis of patients; S.M. contributed to the clinical characterization of patients and the generation of iPSC lines; S.B.R., F.B., R.R., B.X. contributed to scRNAseq data analysis; S.B.R, Z.S., Y.C., R.T, B.X., Y.S., H.Z. contributed to calcium imaging, cell proliferation imaging analysis, PIP-FUCCI biosensor assays and FACS-based neurogenesis analysis; S.B.R, Y.S. and H.Z. contributed to dendritic morphology analysis; K.W.L. supervised the optimization of the organoid culture conditions; S.B.R., B.X. and J.A.G prepared the manuscript with contributions from all co-authors.

## Acknowledgements

This work was supported by the Stavros Niarchos Foundation (SNF), the National Institute of Health-NCATS grant (UG3/UH3TR002151) and National Institute of Mental Health Grant (2R01MH097879). We thank Barbara Corneo and the Columbia Stem Cell Core Facility for the Q1, Q2, Q5 and Q6 hiPSC lines generation. Bio- samples of Q20 and Q27 hiPSCs were obtained from NIMH Repository & Genomics Resource. We thank Linda Brzustowicz and Bill Manley from the Rutgers University and the staff members at RUCDR. Data and biomaterials generated in Study 125/Site 393 were funded by a NIMH grant to Dr. Herb Lachman (MH087840: Analysis of Glutamatergic Neurons Derived from Patient-Specific iPS Cells). The co-investigators on this grant included Dr. Deyou Zheng and Dr. Reed Carroll from the Albert Einstein College of Medicine. Patients and controls were recruited at the Albert Einstein College of Medicine and at the Child Psychiatry Branch, NIMH, directed by Dr. Judith L. Rapoport. We thank all participating subjects and their families for their contributions. We thank the Zuckerman Institute Cellular Imaging core for support with imaging and analysis pipelines for 3D imaging of the organoids and the Zuckerman Institute Flow Cytometry Core for support for FACs sorting. We thank Columbia University Genome Center for support with bulk and scRNAseq execution. We thank Anqi Wang and other members in the Rabadan lab for support with cloud computing and data storage, as well as advice on scRNAseq data analysis.

## Data availability

The raw sequencing data described in this manuscript have been deposited into the NIH Gene Expression Omnibus (GEO) database under accession number GSE244010. The data supporting the findings of this study are available upon request from the corresponding authors.

## SUPPLEMENTAL INFORMATION

## Supplemental Figure

**Fig S1:**
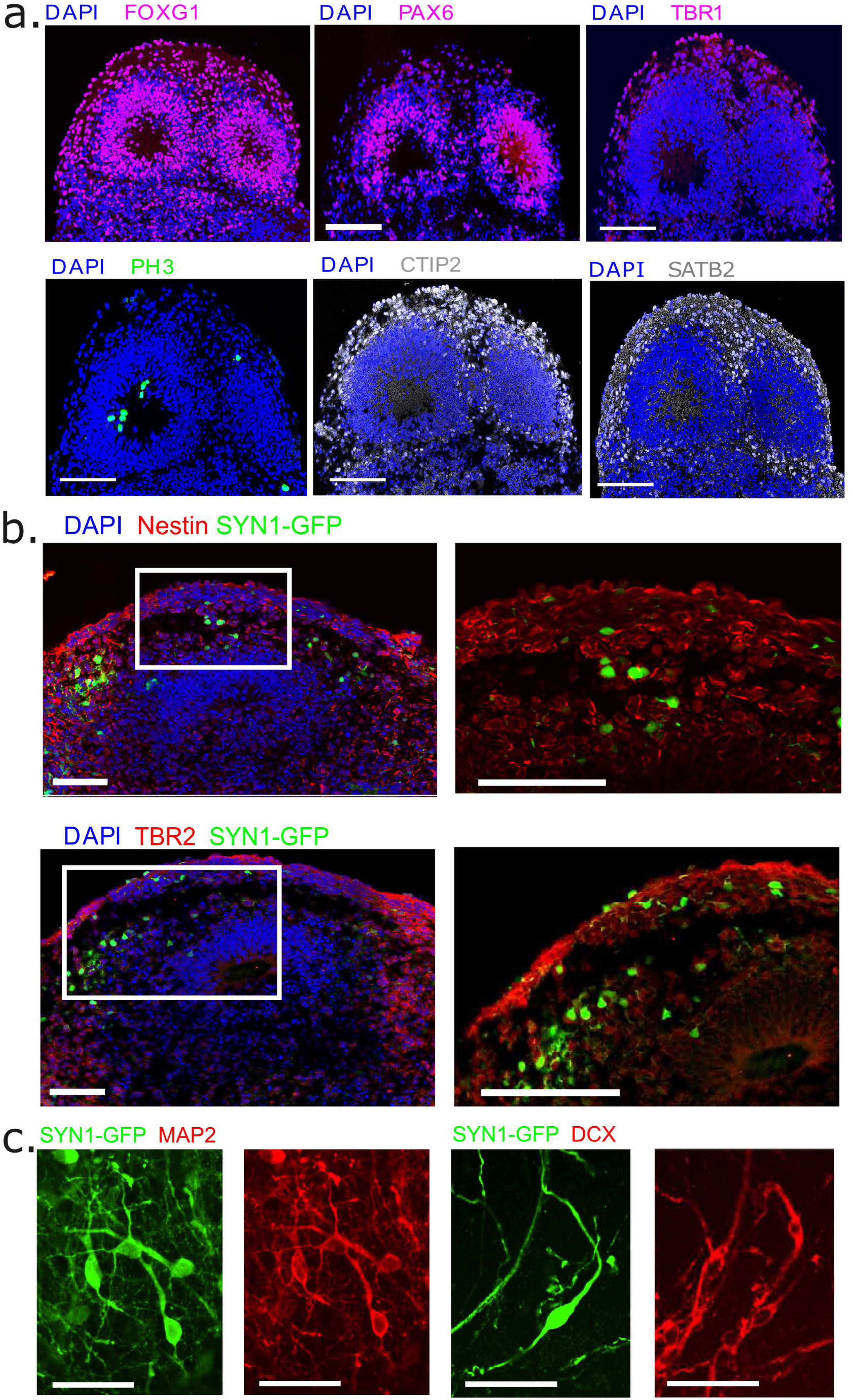
Cell type composition of cultured organoids. Representative confocal images (40x) of 30µm sections from DIV70 control organoids (Control-QR19) immunohistochemically labelled for cell type specific and cell cycle markers: (**a)**. Region specific markers (FOXG1, PAX6, TBR1) and layer specific markers (PH3, CTIP2, and SATB2). Scale bar = 100µm. (**b).** Cell-type specific markers for radial glia (Nestin, *top*) and intermediate progenitor cells (TBR2, *middle*). Scale bar = 50 µm **(c)** Immunofluorescence staining for somatodendritic marker MAP2 (left) and immature neuron marker doublecortin (DCX) (*right*). Scale bar = 20µm. Neurons are labeled with AAV1-hSYN1-GFP.

**Fig S2:**
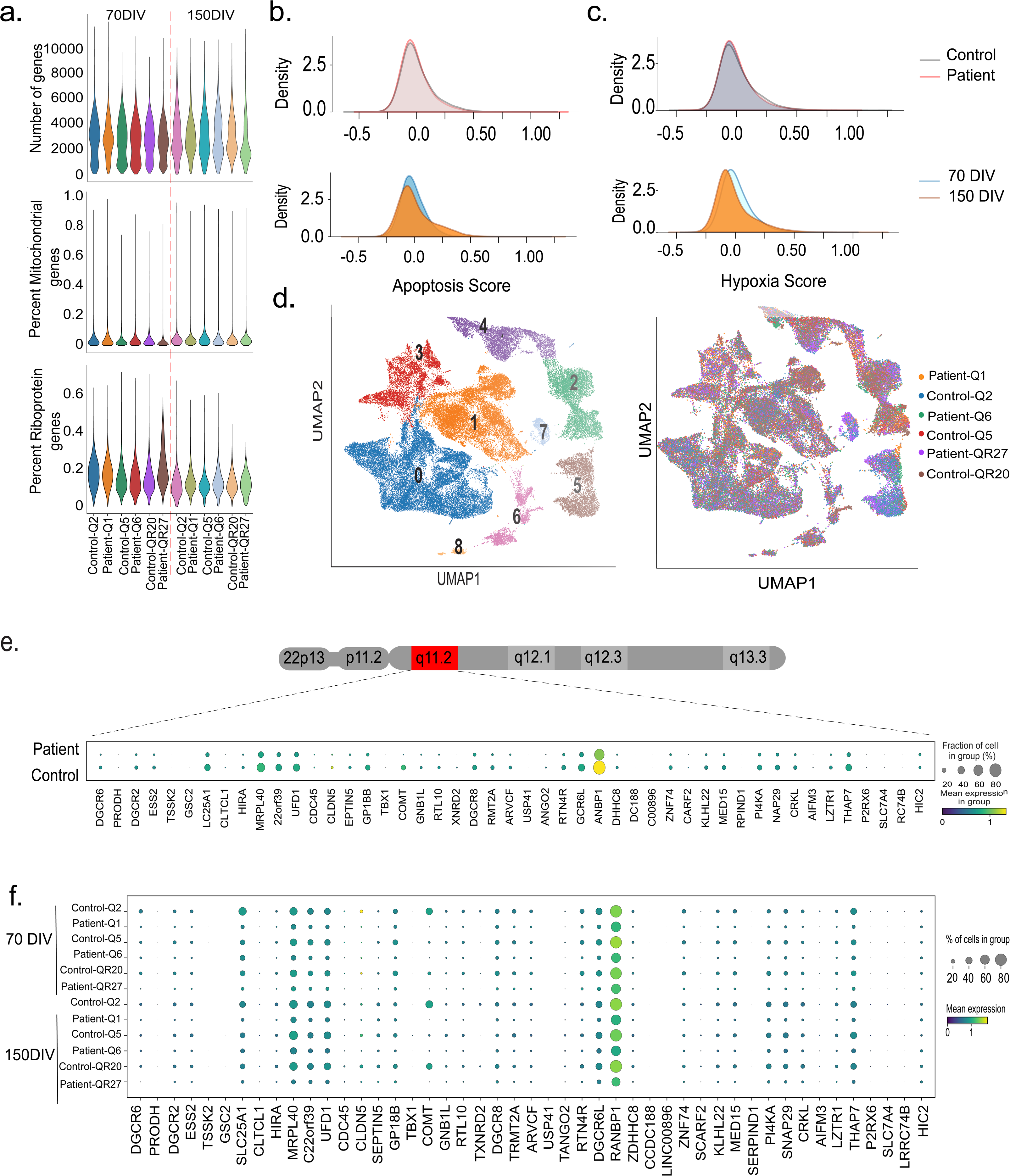
Quality control and clustering metrics of scRNA-seq data. **(a)** Distribution of total number of genes (*top*), mitochondrial transcripts (*middle*) and ribosomal transcripts (*bottom*) per sample. Quality control metrics are uniform across organoids from all iPSC lines. **(b, c)** Expression of genes related to hypoxia and apoptosis curated in the GSEA Hallmark mSigDB database^123^. There is no difference in expression of apoptosis or hypoxia-related genes between patient and control organoids as well as between DIV70 and DIV150 organoids. **(d)** Spatial representation of 8 transcriptomically distinct clusters (*left*) and distribution of clusters in individual samples (*right*). All samples show a similar distribution of clusters. **(e)** Dot-plot depicting expression of 22q11.2 locus genes in scRNA-seq datasets from patient and control organoid. **(f)** Expression of 22q11.2 locus genes across all samples. Analysis confirmed the expected ∼50% decrease in patient relative to control organoids and demonstrated that they faithfully recapitulate the expected 22q11.2 locus gene expression profile with high organoid-to-organoid reproducibility.

**Fig S3:**
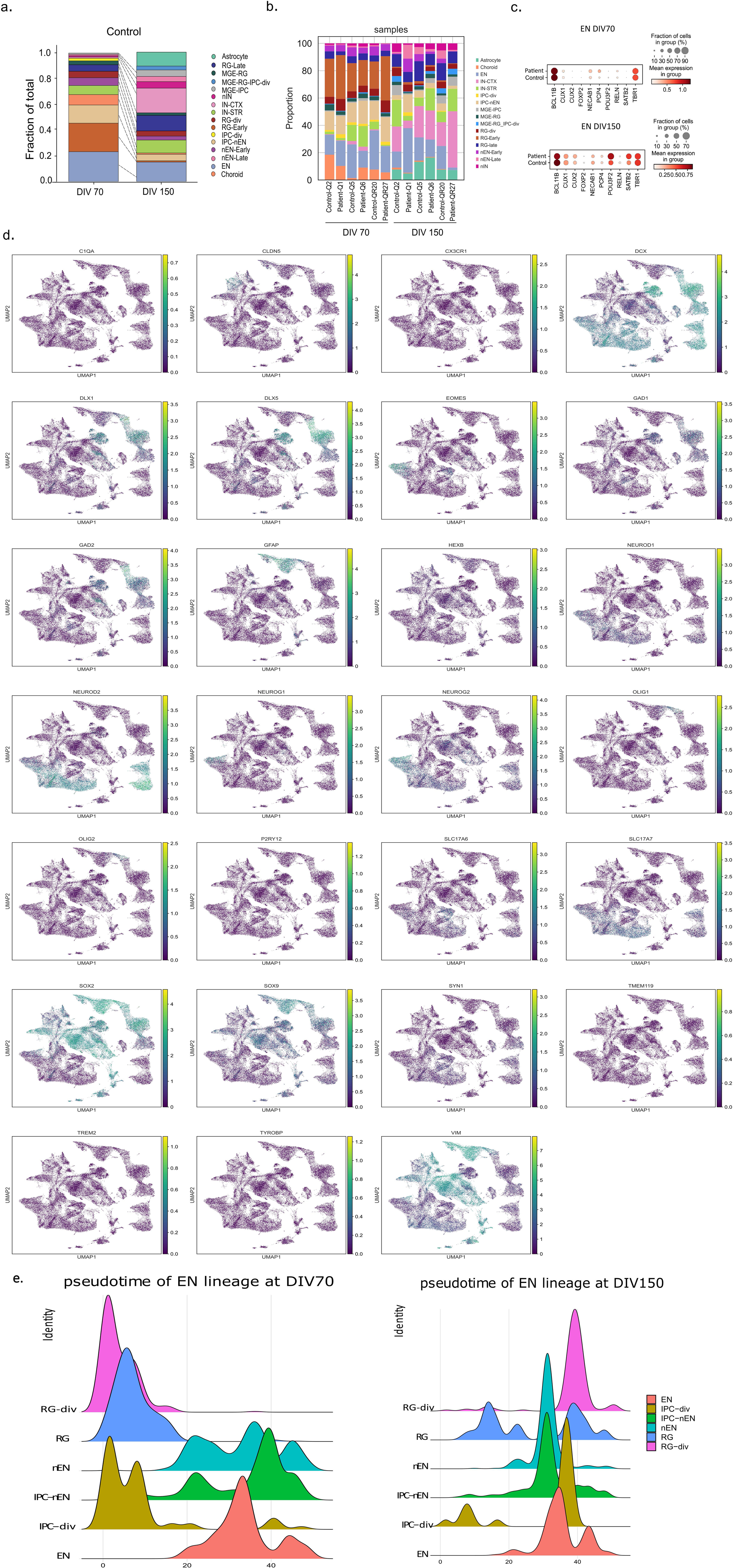
Relative proportion of cell types in control organoids. **(a)** Stacked bar plots depicting the relative proportion of cell types in organoids profiled at DIV70 and DIV150. Early radial glia (RG-early, 22%) and differentiated ENs (23%) are the predominant cell types. Intermediate progenitors differentiating into new born ENs represent the next largest group (IPC-nEN, 14%). 5%-10% of the cell population consists of other intermediate cell types, including dividing radial glia (RG-div), early new born neurons (nEN-early), late radial glia (RG-late), striatal interneurons (IN-STR) and choroid. A minority (≤5%) of cells are comprised of late newborn neurons (nEN-late), cortical interneurons (IN-CTX), dividing intermediate progenitors (IPC-div), MGE progenitors (MGE-RG, MGE-IPC, MGE-RG_IPC-div) and new born interneurons (nIN). Almost no astroglia is detected at DIV70. By DIV150 organoids display a more diverse distribution of cell types including EN (15.5%), RG-late (12%), cortical interneurons (IN-CTX, 19%), IN-STR (10.5%) as well as astrocytes (9.5%). Other cell types represent less than 5% each. The fraction of cells that are part of the EN lineage decreases from 73% at DIV70 to 33% at DIV150 although the proportion of newborn and maturing EN remains fairly constant (30% at DIV70 vs 23% at DIV150). There is an expansion of IN lineage (from 19% at DIV70 to 56% at DIV150), which includes increase in the proportion of IN progenitors (9% to 21%), as well as newborn and maturing IN (10% to 35%). The relative proportion of choroid cells decreases from 7.7% to 0.8% while the astrocyte proportion increases from 0.1% to 9.5%. **(b)** Stacked bar plots depicting the relative proportion of cell types in organoids profiled at DIV70 and DIV150 across all lines. **(c)** Comparison of the mean expression levels of known biomarkers of excitatory neurons (EN) and their expression in fraction of cells in patient and control groups. **(d)** UMAP distribution of various cell type specific markers in organoids. **(e)** Distribution of each cell type in the EN lineage along pseudotime at DIV70 (left panel) and DIV150 in control (right panel).

**Fig S4:**
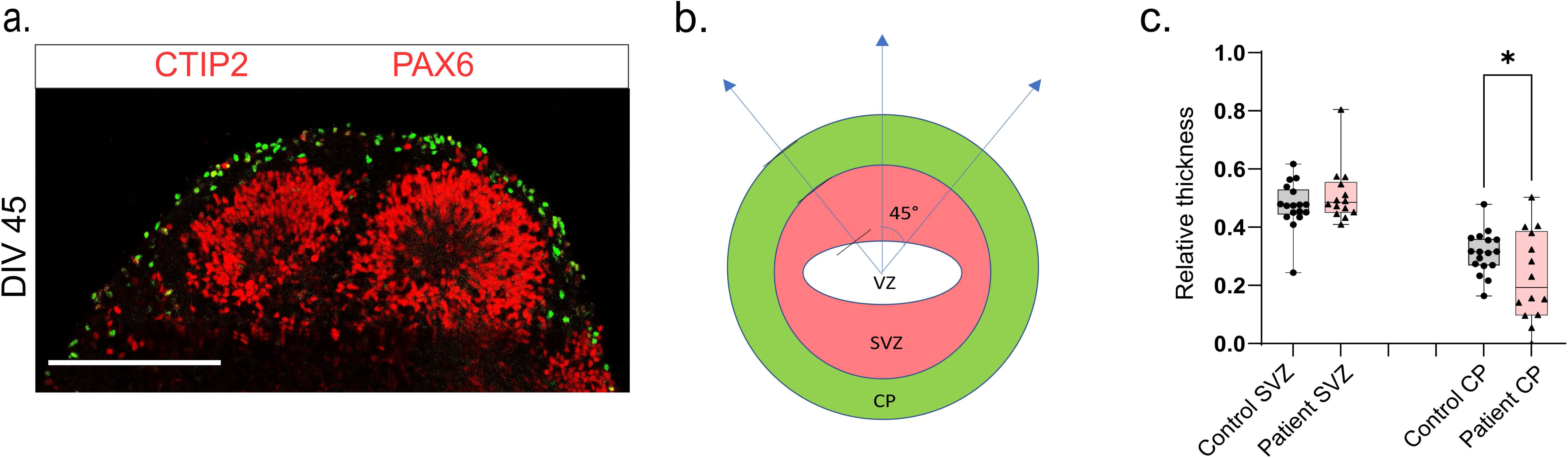
Quantification of the thickness of neurogenic zones of organoids. **(a)** Representative confocal image (40x) of a rosette structure in 30µm section of DIV45 organoids immunostained for PAX6 (red) and CTIP2 (green). Scale bar = 100µm. **(b)** Schematic depicting the quantification of the layer thickness in the rosettes. PAX6 layer is defined as the ventricular zone (VZ), CTIP2 layer is defined as the cortical plate (CP) and total neural tube radius is defined as the distance from the center of the ventricle to the top of the CP layer. Measurements of thickness for each layer were made along 3 lines per rosette that are at 45 degrees to each other and average values for relative thickness of each layer compared to the radius of the rosette (neural tube radius) was calculated. **(c)** Quantification of layer thickness in DIV45 organoids. CP layer thickness is significantly decreased in patient organoids (KS test, *P* = 0.04; total structures analyzed: n ROIs for control = 16, n for patient = 12 from 3 control and 3 patient lines).

**Fig S5:**
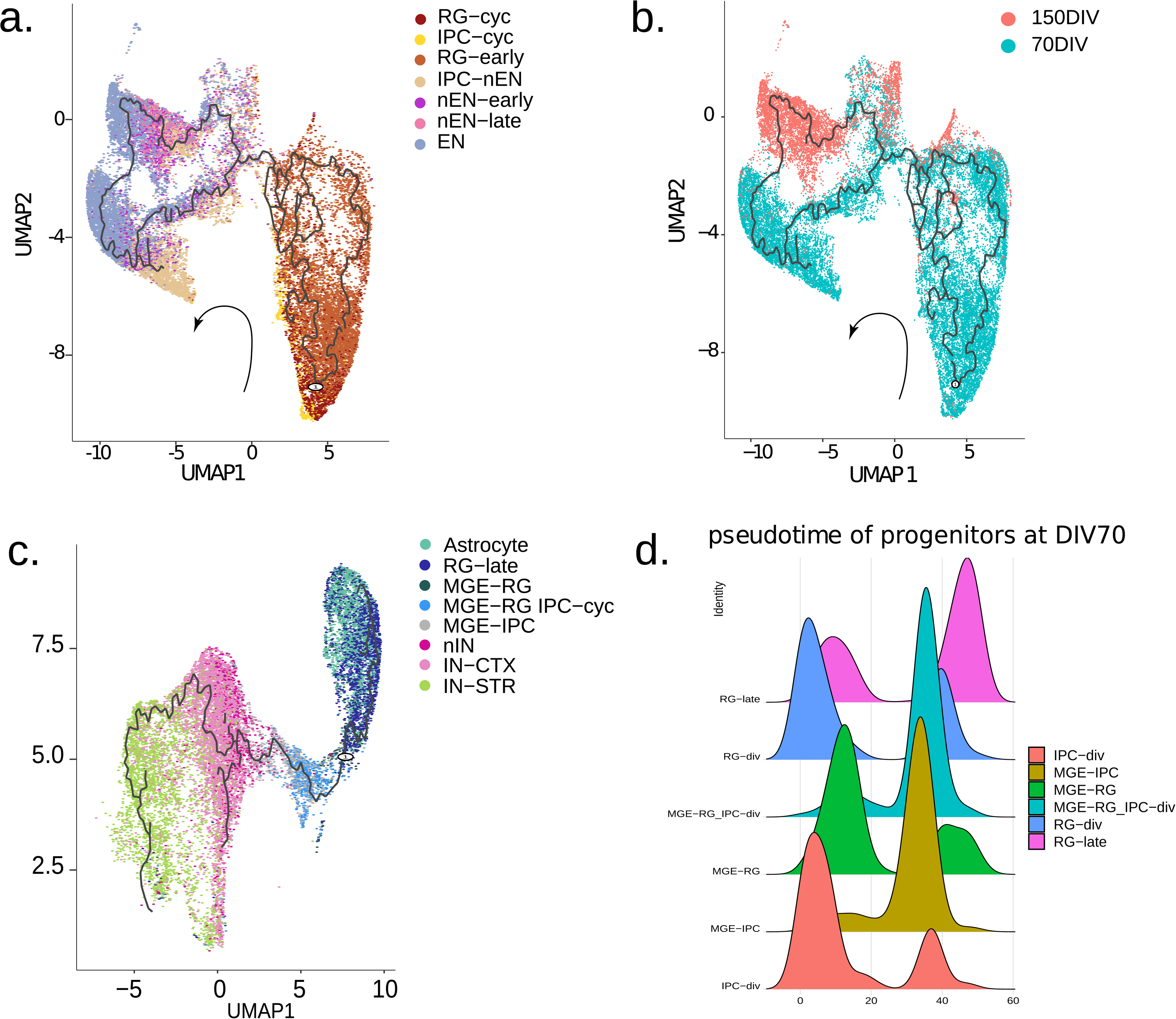
Pseudotime analysis of the EN and IN lineages. **(a)** Pseudotime trajectory of EN lineage calculated with Monocle3 with cell type annotations. Trajectory starts at RG-div and progresses to RG-early, IPC-nEN, then branches into nEN-early and nEN-late and finally to EN. **(b)** Pseudotime trajectory colored by DIV. At DIV70 there are mainly progenitors and neurons in the nEN-early to EN branch. At DIV150 there are fewer early progenitors and mainly neurons in the nEN-late to EN branch. **(c)** Pseudotime trajectory of IN lineage calculated with Monocle3 with cell type annotations. Trajectory starts at RG-late and progresses to MGE RG and MGE IPC, differentiates to nIN, then branches into IN-CTX and IN-STR

**Fig S6:**
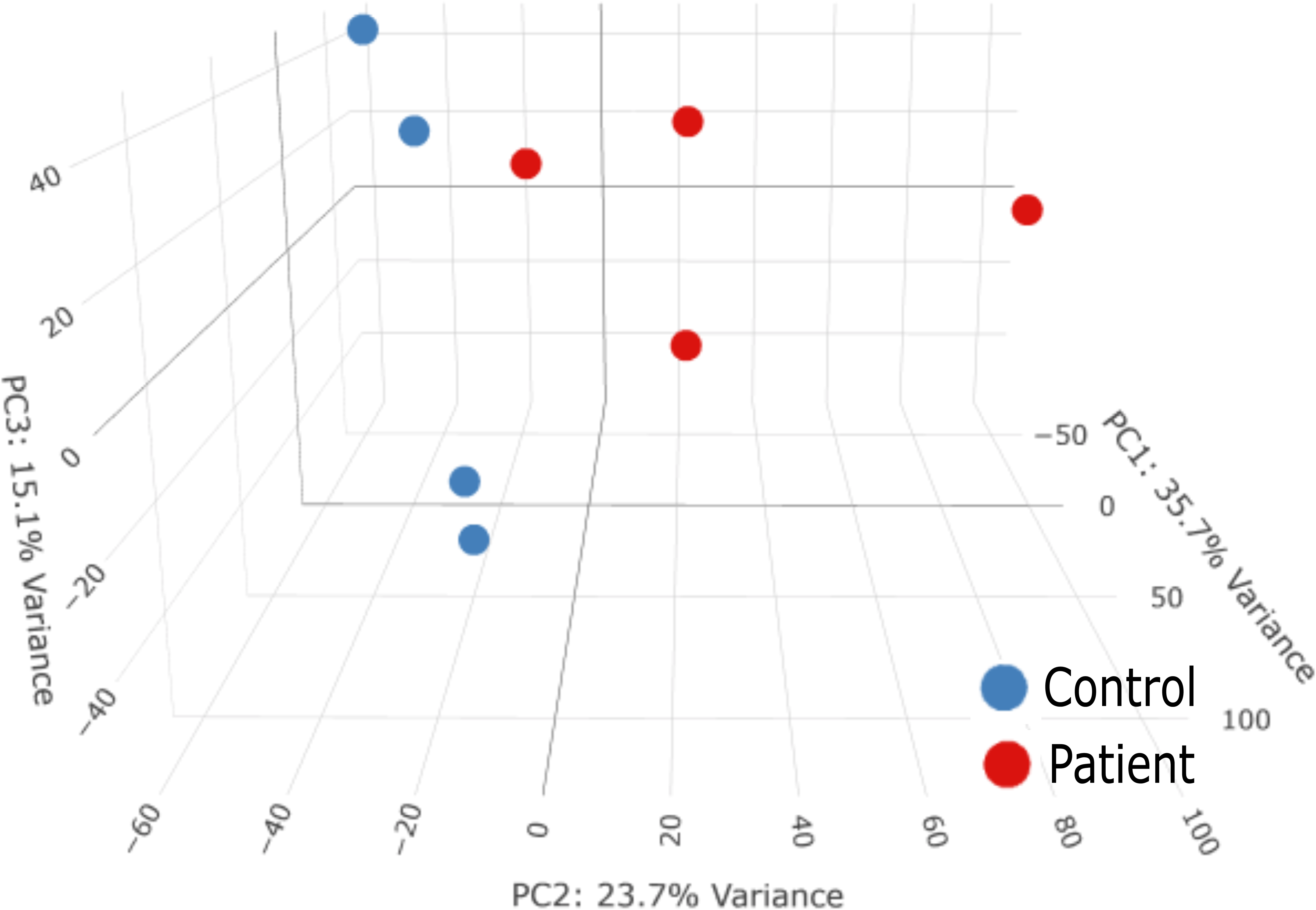
Principle component analysis of gene expression profile in the cortical neurons. The first principal component (PC1) explains 35.7% of the variance that was related to genetic background. PC2 (23.7%) was related to differences between the 22q11DS and control groups.

## Supplemental Tables

**Supplemental Table 1:** Bulk RNAseq_DEGs in in sorted neurons from DIV70 patient organoids compared to control.

**Supplemental Table 2:** scRNAseq_DEGs in various cell types of DIV70 patient organoids compared to control.

**Supplemental Table 3:** scRNAseq_Functional enrichment of DEGs in various cell types of DIV70 patient organoids compared to control.

**Supplemental Table 4:** scRNAseq_DEGs in various cell types of DIV150 patient organoids compared to control.

**Supplemental Table 5:** scRNAseq_Functional enrichment of DEGs in various cell types of DIV150 patient organoids compared to control.

**Supplemental Table 6:** scRNAseq_Functional enrichment of transcription factor regulon targets dysregulated in ENs of DIV70 patient organoids.

**Supplemental Table 7:** WGCNA EN lineage module genes and GO terms.

**Supplemental Table 8:** miRNAseq_Differentially expressed miRNAs in whole patient organoids at DIV70 compared to control.

**Supplemental Table 9:** Clinical and demographic characteristics

**Supplemental Table 10:** Sample usage in experiments

## References

1 Shprintzen, R. J., et al. Velo-cardio-facial syndrome. Curr Opin Pediatr 17, 725–730 (2005). 10.1097/01.mop.0000184465.73833.0b

2 Robin, N. H. & Shprintzen, R. J. Defining the clinical spectrum of deletion 22q11.2. J Pediatr 147, 90–96 (2005). 10.1016/j.jpeds.2005.03.007

3 Kobrynski, L. J. & Sullivan, K. E. Velocardiofacial syndrome, DiGeorge syndrome: the chromosome 22q11.2 deletion syndromes. Lancet 370, 1443–1452 (2007). 10.1016/S0140-6736(07)61601-8

4 Botto, L. D., et al. A population-based study of the 22q11.2 deletion: phenotype, incidence, and contribution to major birth defects in the population. Pediatrics 112, 101–107 (2003). 10.1542/peds.112.1.101

5 Morrison, S., et al. Cognitive deficits in childhood, adolescence and adulthood in 22q11.2 deletion syndrome and association with psychopathology. Transl Psychiatry 10, 53 (2020). 10.1038/s41398-020-0736-7

6 Woodin, M., et al. Neuropsychological profile of children and adolescents with the 22q11.2 microdeletion. Genet Med 3, 34–39 (2001). 10.1097/00125817-200101000-00008

7 Xu, B., et al. Strong association of de novo copy number mutations with sporadic schizophrenia. Nat Genet 40, 880–885 (2008). 10.1038/ng.162

8 Karayiorgou, M., Simon, T. J. & Gogos, J. A. 22q11.2 microdeletions: linking DNA structural variation to brain dysfunction and schizophrenia. Nat Rev Neurosci 11, 402–416 (2010). 10.1038/nrn2841

9 Smerconish, S. & Schmitt, J. E. Neuroanatomical Correlates of Cognitive Dysfunction in 22q11.2 Deletion Syndrome. Genes (Basel) 15 (2024). 10.3390/genes15040440

10 Swillen, A., Moss, E. & Duijff, S. Neurodevelopmental outcome in 22q11.2 deletion syndrome and management. Am J Med Genet A 176, 2160–2166 (2018). 10.1002/ajmg.a.38709

11 Birchwood, M., McGorry, P. & Jackson, H. Early intervention in schizophrenia. Br J Psychiatry 170, 2–5 (1997). 10.1192/bjp.170.1.2

12 Lewis, D. A. & Levitt, P. Schizophrenia as a disorder of neurodevelopment. Annu Rev Neurosci 25, 409–432 (2002). 10.1146/annurev.neuro.25.112701.142754

13 Rapoport, J. L., Addington, A. M., Frangou, S. & Psych, M. R. The neurodevelopmental model of schizophrenia: update 2005. Mol Psychiatry 10, 434–449 (2005). 10.1038/sj.mp.4001642

14 Pantelis, C., et al. Neuroanatomical abnormalities before and after onset of psychosis: a cross-sectional and longitudinal MRI comparison. Lancet 361, 281–288 (2003). 10.1016/S0140-6736(03)12323-9

15 Di Lullo, E. & Kriegstein, A. R. The use of brain organoids to investigate neural development and disease. Nat Rev Neurosci 18, 573–584 (2017). 10.1038/nrn.2017.107

16 Chiaradia, I. & Lancaster, M. A. Brain organoids for the study of human neurobiology at the interface of in vitro and in vivo. Nat Neurosci 23, 1496–1508 (2020). 10.1038/s41593-020-00730-3

17 Velasco, S., Paulsen, B. & Arlotta, P. 3D Brain Organoids: Studying Brain Development and Disease Outside the Embryo. Annu Rev Neurosci 43, 375–389 (2020). 10.1146/annurev-neuro-070918-050154

18 Sun, Z., Williams, D. J., Xu, B. & Gogos, J. A. Altered function and maturation of primary cortical neurons from a 22q11.2 deletion mouse model of schizophrenia. Transl Psychiatry 8, 85 (2018). 10.1038/s41398-018-0132-8

19 Mukai, J., et al. Palmitoylation-dependent neurodevelopmental deficits in a mouse model of 22q11 microdeletion. Nat Neurosci 11, 1302–1310 (2008). 10.1038/nn.2204

20 Stark, K. L., et al. Altered brain microRNA biogenesis contributes to phenotypic deficits in a 22q11-deletion mouse model. Nat Genet 40, 751–760 (2008). 10.1038/ng.138

21 Kadoshima, T., et al. Self-organization of axial polarity, inside-out layer pattern, and species-specific progenitor dynamics in human ES cell-derived neocortex. Proc Natl Acad Sci U S A 110, 20284–20289 (2013). 10.1073/pnas.1315710110

22 Thakur, P. L., Martin; Diamantopoulou, Anastasia ; Rao, Sneha; Chen, Yijing; Ferng, Annie; Mazur, Curt; Kordasiewicz, Holly; Shprintzen, Robert J; Markx, Sander ; Xu, Bin; Gogos, Joseph A. EMC10 reduction in human neurons and adult mouse brain rescues cellular and behavioral deficits linked to 22q11. 2 deletion. biorxiv (2021).

23 Khan, T. A., et al. Neuronal defects in a human cellular model of 22q11.2 deletion syndrome. Nat Med 26, 1888–1898 (2020). 10.1038/s41591-020-1043-9

24 Ilicic, T., et al. Classification of low quality cells from single-cell RNA-seq data. Genome Biol 17, 29 (2016). 10.1186/s13059-016-0888-1

25 Traag, V. A., Waltman, L. & van Eck, N. J. From Louvain to Leiden: guaranteeing well-connected communities. Sci Rep 9, 5233 (2019). 10.1038/s41598-019-41695-z

26 Nowakowski, T. J., et al. Spatiotemporal gene expression trajectories reveal developmental hierarchies of the human cortex. Science 358, 1318–1323 (2017). 10.1126/science.aap8809

27 Aran, D., et al. Reference-based analysis of lung single-cell sequencing reveals a transitional profibrotic macrophage. Nat Immunol 20, 163–172 (2019). 10.1038/s41590-018-0276-y

28 Dann, E., Henderson, N. C., Teichmann, S. A., Morgan, M. D. & Marioni, J. C. Differential abundance testing on single-cell data using k-nearest neighbor graphs. Nat Biotechnol 40, 245–253 (2022). 10.1038/s41587-021-01033-z

29 Qiu, X., et al. Reversed graph embedding resolves complex single-cell trajectories. Nat Methods 14, 979–982 (2017). 10.1038/nmeth.4402

30 Lange, M., et al. CellRank for directed single-cell fate mapping. Nat Methods 19, 159–170 (2022). 10.1038/s41592-021-01346-6

31 Bergen, V., Lange, M., Peidli, S., Wolf, F. A. & Theis, F. J. Generalizing RNA velocity to transient cell states through dynamical modeling. Nat Biotechnol 38, 1408–1414 (2020). 10.1038/s41587-020-0591-3

32 Koopmans, F., et al. SynGO: An Evidence-Based, Expert-Curated Knowledge Base for the Synapse. Neuron 103, 217–234 e214 (2019). 10.1016/j.neuron.2019.05.002

33 Finak, G., et al. MAST: a flexible statistical framework for assessing transcriptional changes and characterizing heterogeneity in single-cell RNA sequencing data. Genome Biol 16, 278 (2015). 10.1186/s13059-015-0844-5

34 Aibar, S., et al. SCENIC: single-cell regulatory network inference and clustering. Nat Methods 14, 1083–1086 (2017). 10.1038/nmeth.4463

35 Feregrino, C. & Tschopp, P. Assessing evolutionary and developmental transcriptome dynamics in homologous cell types. Dev Dyn (2021). 10.1002/dvdy.384

36 Fenelon, K., et al. Deficiency of Dgcr8, a gene disrupted by the 22q11.2 microdeletion, results in altered short-term plasticity in the prefrontal cortex. Proc Natl Acad Sci U S A 108, 4447–4452 (2011). 10.1073/pnas.1101219108

37 Chang, L., Zhou, G., Soufan, O. & Xia, J. miRNet 2.0: network-based visual analytics for miRNA functional analysis and systems biology. Nucleic Acids Res 48, W244–W251 (2020). 10.1093/nar/gkaa467

38 Xu, B., Hsu, P. K., Stark, K. L., Karayiorgou, M. & Gogos, J. A. Derepression of a neuronal inhibitor due to miRNA dysregulation in a schizophrenia-related microdeletion. Cell 152, 262–275 (2013). 10.1016/j.cell.2012.11.052

39 Diamantopoulou, A., et al. Loss-of-function mutation in Mirta22/Emc10 rescues specific schizophrenia-related phenotypes in a mouse model of the 22q11.2 deletion. Proc Natl Acad Sci U S A 114, E6127–E6136 (2017). 10.1073/pnas.1615719114

40 Besson, V., et al. PW1 gene/paternally expressed gene 3 (PW1/Peg3) identifies multiple adult stem and progenitor cell populations. Proc Natl Acad Sci U S A 108, 11470–11475 (2011). 10.1073/pnas.1103873108

41 Kuwahara, A., et al. Tcf3 represses Wnt-beta-catenin signaling and maintains neural stem cell population during neocortical development. PLoS One 9, e94408 (2014). 10.1371/journal.pone.0094408

42 Gribble, S. L., Kim, H. S., Bonner, J., Wang, X. & Dorsky, R. I. Tcf3 inhibits spinal cord neurogenesis by regulating sox4a expression. Development 136, 781–789 (2009). 10.1242/dev.027995

43 Hoshiba, Y., et al. Sox11 Balances Dendritic Morphogenesis with Neuronal Migration in the Developing Cerebral Cortex. J Neurosci 36, 5775–5784 (2016). 10.1523/JNEUROSCI.3250-15.2016

44 D’Rozario, M., et al. Type I bHLH Proteins Daughterless and Tcf4 Restrict Neurite Branching and Synapse Formation by Repressing Neurexin in Postmitotic Neurons. Cell Rep 15, 386–397 (2016). 10.1016/j.celrep.2016.03.034

45 Szklarczyk, D., et al. The STRING database in 2023: protein-protein association networks and functional enrichment analyses for any sequenced genome of interest. Nucleic Acids Res 51, D638–D646 (2023). 10.1093/nar/gkac1000

46 Mencarelli, C., et al. RanBP1 Couples Nuclear Export and Golgi Regulation through LKB1 to Promote Cortical Neuron Polarity. Cell Rep 24, 2529–2539 e2524 (2018). 10.1016/j.celrep.2018.07.107

47 Li, Y. & Jiao, J. Histone chaperone HIRA regulates neural progenitor cell proliferation and neurogenesis via beta-catenin. J Cell Biol 216, 1975–1992 (2017). 10.1083/jcb.201610014

48 Bojjireddy, N., et al. Pharmacological and genetic targeting of the PI4KA enzyme reveals its important role in maintaining plasma membrane phosphatidylinositol 4-phosphate and phosphatidylinositol 4,5-bisphosphate levels. J Biol Chem 289, 6120–6132 (2014). 10.1074/jbc.M113.531426

49 Walsh, R. M., et al. Generation of human cerebral organoids with a structured outer subventricular zone. Cell Rep 43, 114031 (2024). 10.1016/j.celrep.2024.114031

50 Mori, T., et al. Neuroradiological and neurofunctional examinations for patients with 22q11.2 deletion. Neuropediatrics 42, 215–221 (2011). 10.1055/s-0031-1295479

51 Andrews, M. G., et al. LIF signaling regulates outer radial glial to interneuron fate during human cortical development. Cell Stem Cell 30, 1382–1391.e1385 (2023). 10.1016/j.stem.2023.08.009

52 Wang, L., et al. Molecular and cellular dynamics of the developing human neocortex at single-cell resolution. bioRxiv (2024). 10.1101/2024.01.16.575956

53 Velasco, S., et al. Individual brain organoids reproducibly form cell diversity of the human cerebral cortex. Nature 570, 523–527 (2019). 10.1038/s41586-019-1289-x

54 Delgado, R. N., et al. Individual human cortical progenitors can produce excitatory and inhibitory neurons. Nature 601, 397–403 (2022). 10.1038/s41586-021-04230-7

55 Wonders, C. P. & Anderson, S. A. The origin and specification of cortical interneurons. Nature Reviews Neuroscience 7, 687–696 (2006). 10.1038/nrn1954

56 Jorstad, N. L., et al. Transcriptomic cytoarchitecture reveals principles of human neocortex organization. Science 382, eadf6812 (2023). doi:10.1126/science.adf6812

57 Walsh, R. M., et al. Cortical assembloids support the development of fast-spiking human PVALB+ cortical interneurons and uncover schizophrenia-associated defects. bioRxiv (2024). 10.1101/2024.11.26.624368

58 Thapar, A. & Riglin, L. The importance of a developmental perspective in Psychiatry: what do recent genetic-epidemiological findings show? Mol Psychiatry 25, 1631–1639 (2020). 10.1038/s41380-020-0648-1

59 Lek, M., et al. Analysis of protein-coding genetic variation in 60,706 humans. Nature 536, 285–291 (2016). 10.1038/nature19057

60 Burrows, C. K., et al. Genetic Variation, Not Cell Type of Origin, Underlies the Majority of Identifiable Regulatory Differences in iPSCs. PLoS Genet 12, e1005793 (2016). 10.1371/journal.pgen.1005793

61 Kilpinen, H., et al. Common genetic variation drives molecular heterogeneity in human iPSCs. Nature 546, 370–375 (2017). 10.1038/nature22403

62 Kyttälä, A., et al. Genetic Variability Overrides the Impact of Parental Cell Type and Determines iPSC Differentiation Potential. Stem Cell Reports 6, 200–212 (2016). 10.1016/j.stemcr.2015.12.009

63 Zhao, D., et al. MicroRNA Profiling of Neurons Generated Using Induced Pluripotent Stem Cells Derived from Patients with Schizophrenia and Schizoaffective Disorder, and 22q11.2 Del. PLoS One 10, e0132387 (2015). 10.1371/journal.pone.0132387

64 Lin, M., et al. Integrative transcriptome network analysis of iPSC-derived neurons from schizophrenia and schizoaffective disorder patients with 22q11.2 deletion. BMC Syst Biol 10, 105 (2016). 10.1186/s12918-016-0366-0

65 Toyoshima, M., et al. Analysis of induced pluripotent stem cells carrying 22q11.2 deletion. Transl Psychiatry 6, e934 (2016). 10.1038/tp.2016.206

66 Li, J., et al. Mitochondrial deficits in human iPSC-derived neurons from patients with 22q11.2 deletion syndrome and schizophrenia. Transl Psychiatry 9, 302 (2019). 10.1038/s41398-019-0643-y

67 Nehme, R., et al. The 22q11.2 region regulates presynaptic gene-products linked to schizophrenia. Nat Commun 13, 3690 (2022). 10.1038/s41467-022-31436-8

68 Ciceri, G., et al. An epigenetic barrier sets the timing of human neuronal maturation. bioRxiv, 2022.2006.2002.490114 (2022). 10.1101/2022.06.02.490114

69 Gudbrandsen, M., et al. Brain morphometry in 22q11.2 deletion syndrome: an exploration of differences in cortical thickness, surface area, and their contribution to cortical volume. Sci Rep 10, 18845 (2020). 10.1038/s41598-020-75811-1

70 Lin, A., et al. Mapping 22q11.2 Gene Dosage Effects on Brain Morphometry. J Neurosci 37, 6183–6199 (2017). 10.1523/JNEUROSCI.3759-16.2017

71 Sun, D., et al. Large-scale mapping of cortical alterations in 22q11.2 deletion syndrome: Convergence with idiopathic psychosis and effects of deletion size. Mol Psychiatry 25, 1822–1834 (2020). 10.1038/s41380-018-0078-5

72 Bagautdinova, J., et al. Altered cortical thickness development in 22q11.2 deletion syndrome and association with psychotic symptoms. Mol Psychiatry 26, 7671–7678 (2021). 10.1038/s41380-021-01209-8

73 Ching, C. R. K., et al. Mapping Subcortical Brain Alterations in 22q11.2 Deletion Syndrome: Effects of Deletion Size and Convergence With Idiopathic Neuropsychiatric Illness. Am J Psychiatry 177, 589–600 (2020). 10.1176/appi.ajp.2019.19060583

74 Kriegstein, A., Noctor, S. & Martinez-Cerdeno, V. Patterns of neural stem and progenitor cell division may underlie evolutionary cortical expansion. Nat Rev Neurosci 7, 883–890 (2006). 10.1038/nrn2008

75 Lukaszewicz, A., et al. The concerted modulation of proliferation and migration contributes to the specification of the cytoarchitecture and dimensions of cortical areas. Cereb Cortex 16 **Suppl 1**, i26–34 (2006). 10.1093/cercor/bhk011

76 Takahashi, T., Goto, T., Miyama, S., Nowakowski, R. S. & Caviness, V. S., Jr. Sequence of neuron origin and neocortical laminar fate: relation to cell cycle of origin in the developing murine cerebral wall. J Neurosci 19, 10357–10371 (1999). 10.1523/JNEUROSCI.19-23-10357.1999

77 Garcia, K. E., Kroenke, C. D. & Bayly, P. V. Mechanics of cortical folding: stress, growth and stability. Philos Trans R Soc Lond B Biol Sci 373 (2018). 10.1098/rstb.2017.0321

78 Fish, J. L., Dehay, C., Kennedy, H. & Huttner, W. B. Making bigger brains-the evolution of neural-progenitor-cell division. J Cell Sci 121, 2783–2793 (2008). 10.1242/jcs.023465

79 Thakur, P. et al. (eLife Sciences Publications, Ltd, 2024).

80 Ambros, V. MicroRNAs and developmental timing. Curr Opin Genet Dev 21, 511–517 (2011). 10.1016/j.gde.2011.04.003

81 Pasquinelli, A. E. & Ruvkun, G. Control of developmental timing by micrornas and their targets. Annu Rev Cell Dev Biol 18, 495–513 (2002). 10.1146/annurev.cellbio.18.012502.105832

82 La Torre, A., Georgi, S. & Reh, T. A. Conserved microRNA pathway regulates developmental timing of retinal neurogenesis. Proc Natl Acad Sci U S A 110, E2362–2370 (2013). 10.1073/pnas.1301837110

83 Colantuoni, C., et al. Temporal dynamics and genetic control of transcription in the human prefrontal cortex. Nature 478, 519–523 (2011). 10.1038/nature10524

84 Jensen, M. & Girirajan, S. An interaction-based model for neuropsychiatric features of copy-number variants. PLoS Genet 15, e1007879 (2019). 10.1371/journal.pgen.1007879

85 Sanders, S. J., et al. Insights into Autism Spectrum Disorder Genomic Architecture and Biology from 71 Risk Loci. Neuron 87, 1215–1233 (2015). 10.1016/j.neuron.2015.09.016

86 Santinha, A. J., et al. Transcriptional linkage analysis with in vivo AAV-Perturb-seq. Nature 622, 367–375 (2023). 10.1038/s41586-023-06570-y

87 Arstikaitis, P., et al. Paralemmin-1, a modulator of filopodia induction is required for spine maturation. Mol Biol Cell 19, 2026–2038 (2008). 10.1091/mbc.e07-08-0802

88 Mukai, J., et al. Evidence that the gene encoding ZDHHC8 contributes to the risk of schizophrenia. Nat Genet 36, 725–731 (2004). 10.1038/ng1375

89 Stahl, A., et al. Patient iPSC-derived neural progenitor cells display aberrant cell cycle control, p53, and DNA damage response protein expression in schizophrenia. BMC Psychiatry 24, 757 (2024). 10.1186/s12888-024-06127-x

90 Notaras, M., et al. Schizophrenia is defined by cell-specific neuropathology and multiple neurodevelopmental mechanisms in patient-derived cerebral organoids. Molecular Psychiatry 27, 1416–1434 (2022). 10.1038/s41380-021-01316-6

91 Brennand, K. J., et al. Modelling schizophrenia using human induced pluripotent stem cells. Nature 473, 221–225 (2011). 10.1038/nature09915

92 Stern, S., et al. Monozygotic twins discordant for schizophrenia differ in maturation and synaptic transmission. Molecular Psychiatry 29, 3208–3222 (2024). 10.1038/s41380-024-02561-1

93 Paulsen, B., et al. Autism genes converge on asynchronous development of shared neuron classes. Nature 602, 268–273 (2022). 10.1038/s41586-021-04358-6

94 Marshall, C. R., et al. Contribution of copy number variants to schizophrenia from a genome-wide study of 41,321 subjects. Nat Genet 49, 27–35 (2017). 10.1038/ng.3725

95 Xu, B., et al. Exome sequencing supports a de novo mutational paradigm for schizophrenia. Nat Genet 43, 864–868 (2011). 10.1038/ng.902

96 Shohat, S., Ben-David, E. & Shifman, S. Varying Intolerance of Gene Pathways to Mutational Classes Explain Genetic Convergence across Neuropsychiatric Disorders. Cell Rep 18, 2217–2227 (2017). 10.1016/j.celrep.2017.02.007

97 Selemon, L. D. & Zecevic, N. Schizophrenia: a tale of two critical periods for prefrontal cortical development. Transl Psychiatry 5, e623 (2015). 10.1038/tp.2015.115

98 Li, Y., et al. Investigation of Neurodevelopmental Deficits of 22 q11.2 Deletion Syndrome with a Patient-iPSC-Derived Blood-Brain Barrier Model. Cells 10 (2021). 10.3390/cells10102576

99 Mayshar, Y., et al. Identification and classification of chromosomal aberrations in human induced pluripotent stem cells. Cell Stem Cell 7, 521–531 (2010). 10.1016/j.stem.2010.07.017

100 Markouli, C., et al. Gain of 20q11.21 in Human Pluripotent Stem Cells Impairs TGF-beta-Dependent Neuroectodermal Commitment. Stem Cell Reports 13, 163–176 (2019). 10.1016/j.stemcr.2019.05.005

101 Thakur, P., et al. EMC10 reduction in human neurons and adult mouse brain rescues cellular and behavioral deficits linked to 22q11.2 deletion. bioRxiv, 2022.2003.2001.482495 (2022). 10.1101/2022.03.01.482495

102 Dobin, A., et al. STAR: ultrafast universal RNA-seq aligner. Bioinformatics 29, 15–21 (2013). 10.1093/bioinformatics/bts635

103 Love, M. I., Huber, W. & Anders, S. Moderated estimation of fold change and dispersion for RNA-seq data with DESeq2. Genome Biol 15, 550 (2014). 10.1186/s13059-014-0550-8

104 Ge, S. X., Son, E. W. & Yao, R. iDEP: an integrated web application for differential expression and pathway analysis of RNA-Seq data. BMC Bioinformatics 19, 534 (2018). 10.1186/s12859-018-2486-6

105 Raudvere, U., et al. g:Profiler: a web server for functional enrichment analysis and conversions of gene lists (2019 update). Nucleic Acids Res 47, W191–W198 (2019). 10.1093/nar/gkz369

106 Fan, Y., et al. miRNet - dissecting miRNA-target interactions and functional associations through network-based visual analysis. Nucleic Acids Res 44, W135–141 (2016). 10.1093/nar/gkw288

107 Gel, B., et al. regioneR: an R/Bioconductor package for the association analysis of genomic regions based on permutation tests. Bioinformatics 32, 289–291 (2016). 10.1093/bioinformatics/btv562

108 Zheng, G. X., et al. Massively parallel digital transcriptional profiling of single cells. Nat Commun 8, 14049 (2017). 10.1038/ncomms14049

109 Wolf, F. A., Angerer, P. & Theis, F. J. SCANPY: large-scale single-cell gene expression data analysis. Genome Biol 19, 15 (2018). 10.1186/s13059-017-1382-0

110 Lun, A. T., Bach, K. & Marioni, J. C. Pooling across cells to normalize single-cell RNA sequencing data with many zero counts. Genome Biol 17, 75 (2016). 10.1186/s13059-016-0947-7

111 Welch, J. D., et al. Single-Cell Multi-omic Integration Compares and Contrasts Features of Brain Cell Identity. Cell 177, 1873–1887 e1817 (2019). 10.1016/j.cell.2019.05.006

112 Becht, E., et al. Dimensionality reduction for visualizing single-cell data using UMAP. Nat Biotechnol (2018). 10.1038/nbt.4314

113 Trapnell, C., et al. The dynamics and regulators of cell fate decisions are revealed by pseudotemporal ordering of single cells. Nat Biotechnol 32, 381–386 (2014). 10.1038/nbt.2859

114 Van de Sande, B., et al. A scalable SCENIC workflow for single-cell gene regulatory network analysis. Nat Protoc 15, 2247–2276 (2020). 10.1038/s41596-020-0336-2

115 Berg, S., et al. ilastik: interactive machine learning for (bio)image analysis. Nat Methods 16, 1226–1232 (2019). 10.1038/s41592-019-0582-9

116 Schindelin, J., et al. Fiji: an open-source platform for biological-image analysis. Nat Methods 9, 676–682 (2012). 10.1038/nmeth.2019

117 Legland, D., Arganda-Carreras, I. & Andrey, P. MorphoLibJ: integrated library and plugins for mathematical morphology with ImageJ. Bioinformatics 32, 3532–3534 (2016). 10.1093/bioinformatics/btw413

118 Grant, G. D., Kedziora, K. M., Limas, J. C., Cook, J. G. & Purvis, J. E. Accurate delineation of cell cycle phase transitions in living cells with PIP-FUCCI. Cell Cycle 17, 2496–2516 (2018). 10.1080/15384101.2018.1547001

119 Hayatigolkhatmi, K. et al. (eLife Sciences Publications, Ltd, 2024).

120 Giovannucci, A., et al. CaImAn an open source tool for scalable calcium imaging data analysis. Elife 8 (2019). 10.7554/eLife.38173

121 Fromer, M., et al. Gene expression elucidates functional impact of polygenic risk for schizophrenia. Nat Neurosci 19, 1442–1453 (2016). 10.1038/nn.4399

122 Ashtiani, M., Mirzaie, M. & Jafari, M. CINNA: an R/CRAN package to decipher Central Informative Nodes in Network Analysis. Bioinformatics 35, 1436–1437 (2019). 10.1093/bioinformatics/bty819

123 Liberzon, A., et al. The Molecular Signatures Database (MSigDB) hallmark gene set collection. Cell Syst 1, 417–425 (2015). 10.1016/j.cels.2015.12.004

